# Coordinated *Tbx3/Tbx5* transcriptional control of the adult ventricular conduction system

**DOI:** 10.1101/2024.08.29.610377

**Authors:** Ozanna Burnicka-Turek, Katy A. Trampel, Brigitte Laforest, Michael T. Broman, Xinan H. Yang, Zoheb Khan, Eric Rytkin, Binjie Li, Ella Schaffer, Margaret Gadek, Kaitlyn M. Shen, Igor R. Efimov, Ivan P. Moskowitz

## Abstract

The cardiac conduction system (CCS) orchestrates the electrical impulses that enable coordinated contraction of the cardiac chambers. The T-box transcription factors *TBX3* and *TBX5* are required for cardiac conduction system development and associated with overlapping and distinct human cardiac conduction system diseases. We evaluated the coordinated role of *Tbx3* and *Tbx5* in the murine ventricular conduction system (VCS). We engineered a compound *Tbx3:Tbx5* conditional knockout allele for both genes located in *cis* on mouse chromosome 5. Conditional deletion of both T-box transcriptional factors in the ventricular conduction system, using the VCS-specific *MinK:Cre,* caused loss of VCS function and molecular identity. Combined *Tbx3* and *Tbx5* deficiency in the adult VCS led to conduction defects, including prolonged PR and QRS intervals and elevated susceptibility to ventricular tachycardia. These electrophysiological defects occurred prior to detectable alterations in cardiac contractility or histologic morphology, indicative of a primary conduction system defect. *Tbx3:Tbx5* double knockout VCS cardiomyocytes revealed a transcriptional shift towards non-CCS-specialized working myocardium, indicating a change to their cellular identity. Furthermore, optical mapping revealed a loss of VCS-specific conduction system propagation. Collectively, these findings indicate that *Tbx3* and *Tbx5* coordinate to control VCS molecular fate and function, with implications for understanding cardiac conduction disorders in humans.

## Introduction

The cardiac conduction system (CCS) constitutes a highly specialized network of cardiomyocytes that initiate and propagate the electrical impulses required for synchronized contractions of the heart. In the mature mammalian heart, the functional components of the CCS can be broadly divided into the slowly propagating atrial nodes (∼ 5cm/sec), containing the sinoatrial node (SAN) and atrioventricular node (AVN), and the rapidly propagating ventricular conduction system (VCS) (∼ 200cm/sec), including the AV (His) bundle (AVB) and the right and left bundle branches (BBs). The VCS is responsible for rapid propagation of the electrical impulse from the AVN to the ventricular apex to enable synchronous ventricular contraction and effective ejection of blood from the ventricles (1–3). Defects of CCS can occur in normally formed hearts as well as in patients with structural congenital heart disease and are a major source of morbidity and mortality (1, 3–6). The VCS specifically has been recognized as a substrate for life-threatening ventricular arrhythmias, including bundle branch reentry tachycardia, idiopathic fascicular tachycardia, short-coupled torsade de pointes, and ventricular fibrillation (1, 6–9). Despite the severe clinical consequences of CCS disorders, the molecular mechanisms that establish and maintain regional functionality of the mature CCS domains require further study.

Human genetic studies have identified numerous loci associated with adult human CCS function, including the developmentally important factors *Tbx3* and *Tbx5* (reviewed in 1, 10). *Tbx3* and *Tbx5* play crucial roles in adult CCS development and function (1, 3, 11–20). *Tbx5* encodes a T-box transcriptional activator required for structural and conduction system cardiac development (3, 12, 20, 21). Dominant mutations in human *TBX5* cause Holt-Oram syndrome (HOS, OMIM:142900), an autosomal dominant disorder characterized by upper limb malformations, congenital heart defects and CCS abnormalities (22–24). The cardiac phenotype of HOS, including atrioventricular conduction delay, has been recapitulated in the *Tbx5* heterozygous mice (20). Moreover, VCS-specific *Tbx5* knockout caused slowed VCS function and ventricular tachycardia resulting in sudden death in mice (1), emphasizing the importance of *Tbx5* in VCS conduction. *Tbx5* is strongly expressed in the atria and VCS (1, 3, 12, 17), and directly regulates several targets required for VCS function (3, 20, 25), including *Gja5 (Cx40)* (20) and *Scn5a (Nav1.5)* (1). *Tbx3* encodes a T-box transcriptional repressor which is critical for cardiac development (17, 26). Dominant mutations in human *TBX3* cause Ulnar-Mammary syndrome (OMIM:181450), a developmental disorder (27, 28), that includes functional conduction system defects (29). In the heart, *Tbx3* is specifically expressed within CCS (17, 30), and its deficiency below critical level leads to lethal arrhythmias (26). Furthermore, *Tbx3* is required for the molecular identity but not the function of the VCS (17). In contrast, *Tbx3* in SAN and AVN is required for their proper function (30), emphasizing its critical role in maintaining proper cardiac rhythm.

A model for regional CCS specialization suggests that the adult CCS is organized entirely as a slow conduction system ground state by *Tbx3* with a T-box-dependent, physiologically dominant fast conduction system network driven specifically in the VCS by *Tbx5* (11). The adult VCS-specific removal of TBX5 or overexpression of TBX3 shifted the fast VCS into a slow nodal-like system, indicating that the *Tbx3/Tbx5* ratio determines nodal versus VCS function (11). However, a comprehensive assessment of the coordinated requirements for *Tbx3* and *Tbx5* has been hindered by the inability to achieve their compound deletion due to their genomic proximity. *Tbx3* and *Tbx5* are situated in *cis* within 0.6 Mb on chromosome 5 in mice, rendering their simultaneous deletion unattainable with the available single allele conditional alleles. To investigate the consequences of *Tbx3* and *Tbx5* compound removal from the mature VCS, we generated a novel compound *Tbx3:Tbx5* double conditional allele. We found that VCS-specific genetic removal of both TBX3 and TBX5 transformed fast-conducting, adult VCS into working myocardium-like cardiomyocytes, shifting them from conduction to non- conduction myocytes. These results demonstrated the coordinated requirements of both *Tbx3* and *Tbx5* for maintained specification of the mature ventricular conduction system.

## Results

We generated a novel *Tbx3:Tbx5* double-floxed allele to enable the simultaneous conditional deletion of *Tbx3* and *Tbx5* genes specifically from the adult VCS. *Tbx3* and *Tbx5* reside in *cis* on mouse chromosome 5 (*Tbx3* mm39 chr5:119808734-119822789; *Tbx5* mm39 chr5:119970733-120023284). Therefore, to generate a double-conditional knockout, we targeted *Tbx5* in the background of a previously validated *Tbx3* floxed allele (26) using the CRISPR-Cas9 system (31, 32) (Figure 1A). We engineered a *Tbx5* floxed allele mirroring a previously published allele (20). This design enabled us to utilize the previously published individual *Tbx3* floxed allele (26) and individual *Tbx5* floxed allele (20) to serve as controls (Figure 1A and Supplementary Figure 1 and 2).

**Figure 1.**
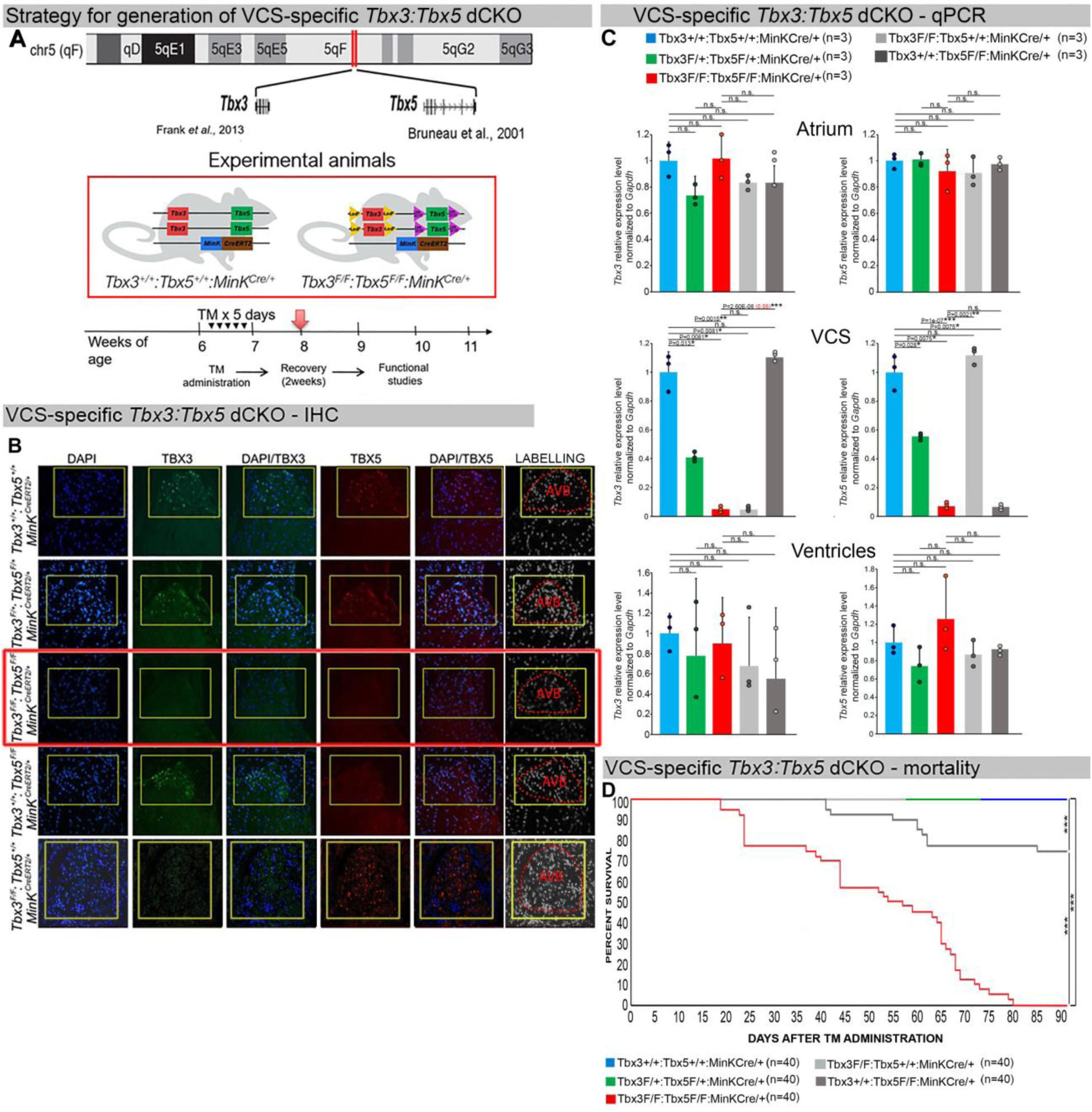
Generation of VCS-specific *Tbx3:Tbx5* double-conditional knockout mice. **(A)** Strategy to generate VCS-specific Tbx3*:Tbx5* double-conditional knockout mouse line. A new *Tbx3:Tbx5* double-conditional knockout mouse line (*Tbx3^fl/fl^;Tbx5^fl/fl^*) was generated using the CRISPR-Cas9 system (31, 32) which allowed for the targeting of *Tbx5* in the background of the previously validated *Tbx3* floxed allele (26). A newly engineered *Tbx5* floxed allele has been developed to mirror a previously published allele (20). This design has enabled the utilization of the previously published individual *Tbx3* floxed allele (26) and individual *Tbx5* floxed allele (20) as controls. To conditionally delete *Tbx3* and *Tbx5* genes specifically from the adult VCS and generate the experimental animals, the *Tbx3:Tbx5* double-floxed mouse line (*Tbx3^fl/fl^;Tbx5^fl^/^fl^*) was combined with a VCS-specific tamoxifen inducible *Cre* transgenic mouse line (*MinK^CreERT2^* [*Tg(RP23- 276I20-MinKCreERT2*) (8). All allelic combinations were generated and evaluated as littermates in a mixed genetic background. The experimental mice employed in all studies were administered tamoxifen at 6 weeks of age and subsequently evaluated at 9 weeks of age (3 weeks post-tamoxifen administration). **(B, C)** The loss of *Tbx3* and *Tbx5* expression, on both the protein and mRNA levels, assessed by immunohistochemistry and qRT-PCR **(C)**, respectively, was observed in the VCS of adult *Tbx3:Tbx5* double- conditional mutant mice (*Tbx3^fl/fl^;Tbx5^fl/fl^;R26^EYFP/+^;MinK^CreERT2/+^)*, but not in their littermate controls (*Tbx3^+/+^;Tbx5^+/+^;R26^EYFP/+^;MinK^CreERT2/+^*) **(B and C)**. **(C)** QRT-PCR analysis showed a partial loss of *Tbx3* and *Tbx5* expression in the adult VCS of *Tbx3:Tbx5* double- conditional heterozygous mice (*Tbx3^fl/+^;Tbx5^fl/+^;R26^EYFP/+^;MinK^CreERT2/+^)* compared to their littermate controls (*Tbx3^+/+^;Tbx5^+/+^;R26^EYFP/+^;MinK^CreERT2/+^*). Additionally, qRT-PCR analysis confirmed the specificity of the *Tbx3:Tbx5* double knockout for the VCS by assessing *Tbx3* and *Tbx5* expression levels in the atria and ventricles of tamoxifen- treated experimental mice. Consistent with the VCS selectivity of *Cre* activity in the *MinKCreERT2* mice (8), *Tbx3* and *Tbx5* expression remained similar in the atrial and ventricular myocardium across all allelic combinations, including *Tbx3:Tbx5* double-conditional knockout mice (*Tbx3^fl/fl^;Tbx5^fl/fl^;R26^EYFP/+^;MinK^CreERT2/+^*). **(D)** Conducted longitudinal studies revealed a significantly increased mortality rate in VCS-specific *Tbx3:Tbx5-*deficient mice compared to their littermate controls (***P<0.0001, log-rank test, GraphPad Prism), suggesting a requirement for both *Tbx3* and *Tbx5* in the mature VCS. All allelic combinations of experimental and control mice (n=40 biological replicates/genotype) were followed longitudinally after tamoxifen administration at 6 weeks of age. *Tbx3:Tbx5* double-conditional knockout mice began to die suddenly at 3 to 4 weeks post-tamoxifen administration. Within the 3 months post-tamoxifen administration, all tamoxifen-treated *Tbx3^fl/fl^;Tbx5^fl/fl^;R26^EYFP/+^;MinK^CreERT2/+^* mice had died suddenly (n=40) without previous signs of illness. In contrast, no mortality was observed among the tamoxifen-treated *Tbx3^+/+^;Tbx5^+/+^;R26^EYFP/+^;MinK^CreERT2/+^ and Tbx3^fl/+^;Tbx5^fl/+^;R26^EYFP/+^;MinK^CreERT2/+^* littermates (each cohort n=40) during this period. TBX3 and TBX5 protein expression was evaluated by immunohistochemistry (green and red signals, respectively) on serial sections from hearts of all allelic combinations (n=3 biological replicates/genotype). Nuclei were stained with DAPI (blue signal). IHC original magnification: 40x. QRT-PCR data are presented as mean±SD normalized to *Gapdh* and relative to *Tbx3^+/+^;Tbx5^+/+^;R26^EYFP/+^;MinK^CreERT2/+^* mice (set as 1). N=3 biological replicates/genotype (VCS cardiomyocytes pooled from 30 mice per biological replicate); multiple testing correction using Benjamini & Hochberg procedure. Significance was assessed by Welch *t* test (*FDR<0.05; **FDR<0.005; *** FDR<0.001) and confirmed by one-tail Wilcoxon test (P value in parentheses) when normally distribution was rejected. Abbreviations: AVB, atrioventricular bundle (also known as bundle of His); FDR, false discovery rate; VCS, ventricular conduction system.

We assessed the impact of removing *Tbx3* and *Tbx5* from the mature VCS by combining the *Tbx3:Tbx5* double-floxed mouse line (*Tbx3^fl/fl^;Tbx5^fl/fl^*) with a VCS-specific tamoxifen (TM) inducible *Cre* transgenic mouse line (MinK^CreERT2^ [Tg(RP23-276I20- *MinKCreERT2)* (Figure 1A) (8). Individual *Tbx3* floxed and *Tbx5* floxed mouse lines combined with *MinK^CreERT2^* transgenic mouse lines (*Tbx3^fl/fl^;Tbx5^+/+^; R26^EYFP/+^;MinK^CreERT2/+^* and *Tbx3^+/+^;Tbx5^fl/fl^;R26^EYFP/+^;MinK^CreERT2/+^,* respectively), were generated as controls, and all allelic combinations were evaluated in a mixed genetic background. We compared VCS-specific *Tbx3:Tbx5* double-conditional mutants (*Tbx3^fl/fl^;Tbx5^fl/fl^;R26^EYFP/+^;MinK^CreERT2/+^)* with control littermates (*Tbx3^+/+^;Tbx5^+/+^; R26^EYFP/+^;MinK^CreERT2/+^*) and VCS-specific *Tbx3:Tbx5* double-conditional heterozygous littermates (*Tbx3^fl/+^;Tbx5^fl/+^;R26^EYFP/+^;MinK^CreERT2/+^).* Additionally, we validated the newly created *Tbx5* floxed allele demonstrating that it is efficiently converted to the *Tbx5* null allele through *Cre* recombinase, causing a phenotype consistent with that observed from conversion of the previously published *Tbx5* floxed allele (1, 20) (Supplementary Figure 1 and 2, Supplementary Material section).

We assessed experimental mice at 8-9 weeks of age following tamoxifen administration at 6 weeks of age (Figure 1A, Methods section). We observed loss of both *Tbx3* and *Tbx5* expression, on both the mRNA and protein levels, in the VCS of adult *Tbx3:Tbx5* double-conditional mutant mice (*Tbx3^fl/fl^;Tbx5^fl/fl^;R26^EYFP/+^;MinK^CreERT2/+^)* but not in their littermate controls (*Tbx3^+/+^;Tbx5^+/+^;R26^EYFP/+^;MinK^CreERT2/+^*) (Figure 1B and C). Partial loss of *Tbx3* and *Tbx5* expression in the adult VCS of *Tbx3:Tbx5* double-conditional heterozygous mice (*Tbx3^fl/+^;Tbx5^fl/+^;R26^EYFP/+^;MinK^CreERT2/+^)* was observed compared to littermate controls (*Tbx3^+/+^;Tbx5^+/+^;R26^EYFP/+^;MinK^CreERT2/+^*) (Figure 1C). We confirmed the specificity of the *Tbx3:Tbx5* double knockout for the VCS by assessing *Tbx3* and *Tbx5* expression levels in the atria and ventricles of tamoxifen-treated experimental mice. Consistent with the VCS selectivity of *Cre* activity in the *MinK^CreERT2^* mice (8), *Tbx3* and *Tbx5* expression remained similar in the atrial and ventricular myocardium of all allelic combinations, including *Tbx3:Tbx5* double-conditional knockout mice (*Tbx3^fl/fl^;Tbx5^fl/fl^;R26^EYFP/+^;MinK^CreERT2/+^*) (Figure 1C).

*Tbx3:Tbx5* double-conditional knockout mice (*Tbx3^fl/fl^;Tbx5^fl/fl^;R26^EYFP/+^; MinK^CreERT2/+^)* appeared morphologically and functionally normal and indistinguishable from control littermates (*Tbx3^+/+^;Tbx5^+/+^;R26^EYFP/+^;MinK^CreERT2/+^*) at 2 weeks post-tamoxifen. However, longitudinal analysis demonstrated sudden death of VCS-specific double knockout mice beginning 3 weeks post-tamoxifen administration (Figure 1D). Within the 3 months post-tamoxifen administration, all tamoxifen-treated *Tbx3^fl/fl^;Tbx5^fl/fl^; R26^EYFP/+^;MinK^CreERT2/+^* mice had died suddenly (n=40). In contrast, no mortality was observed among the tamoxifen-treated *Tbx3^+/+^;Tbx5^+/+^;R26^EYFP/+^;MinK^CreERT2/+^* or *Tbx3^fl/+^;Tbx5^fl/+^;R26^EYFP/+^;MinK^CreERT2/+^* littermates (each cohort n=40) during this period (*Tbx3:Tbx5* double-conditional knockout mice vs control mice P<0.0001; *Tbx3:Tbx5* double-conditional knockout mice vs *Tbx3:Tbx5* double-conditional heterozygous mice P<0.0001, log-rank test; Figure 1D). These results revealed that double deletion of *Tbx3* and *Tbx5* from the adult VCS causes lethality beginning at 3 weeks post-tamoxifen.

The onset of mortality observed in VCS-specific *Tbx3:Tbx5-*deficient mice starting at 3 weeks post-tamoxifen prompted us to investigate the electrophysiologic consequences of VCS-specific *Tbx3:Tbx5* double-knockout at 2-3 weeks post-tamoxifen, prior to the onset of lethality (Figure 2). VCS-specific *Tbx3:Tbx5*-deficiency caused profound conduction slowing in *Tbx3^fl/fl^;Tbx5^fl/fl^;R26^EYFP/+^;MinK^CreERT2/+^* mice by ambulatory telemetry ECG analysis compared to *Tbx3:Tbx5* double-conditional heterozygous mice (*Tbx3^fl/+^;Tbx5^fl/+^;R26^EYFP/+^;MinK^CreERT2/+^*) and littermate controls (*Tbx3^+/+^;Tbx5^+/+^;R26^EYFP/+^;MinK^CreERT2/+^*) (Figure 2A-F). Specifically, the PR interval, representing the period between atrial and ventricular depolarization, and the QRS duration, indicating the length of ventricular depolarization and early repolarization in mice, were both significantly prolonged (Figure 2A, B, and D; PR: *Tbx3:Tbx5* double-conditional knockout mice vs control mice P<0.05, n=7, Welch *t* test; QRS: *Tbx3:Tbx5* double-conditional knockout mice vs control mice P<0.05, n=7, Welch t test). Removal of both *Tbx3* and *Tbx5* from the adult VCS resulted in increased episodes of spontaneous ventricular tachycardia (VT). Ambulatory studies revealed episodes of spontaneous VT in 4 out of 7 *Tbx3^fl/fl^;Tbx5^fl/fl^;R26^EYFP/+^;MinK^CreERT2/+^* mice, in contrast to none observed in 7 littermate controls (Figure 2G, P<0.05, n=7, Welch *t* test). Furthermore, *Tbx3:Tbx5* double-conditional knockout mice (*Tbx3^fl/fl^;Tbx5^fl/fl^;R26^EYFP/+^;MinK^CreERT2/+^)* showed significantly increased susceptibility to ventricular tachycardia following burst stimulation in invasive electrophysiology (EP) studies (3 of 3 *Tbx3^fl/fl^;Tbx5^fl/fl^;R26^EYFP/+^;MinK^CreERT2/+^* mice versus 0 of 7 littermate controls; (Figure 2H, P<0.05, Welch *t* test). In contrast, VCS-specific *Tbx3:Tbx5* double-conditional heterozygous mice (n=7) showed neither conduction nor electrophysiological defects (Figure 2). Consistent with the use of a VCS-specific *Cre* (*MinK-Cre*), no changes in the refractory/recovery periods of atrium, ventricle, or nodes (atrial effective refractory period, ventricular effective refractory period, atrioventricular nodal effective refractory period, or sinus node recovery time) were detected by intracardiac electrophysiology conducted on experimental and control mice (Figure 2I, P<0.05, Welch *t* test).

**Figure 2.**
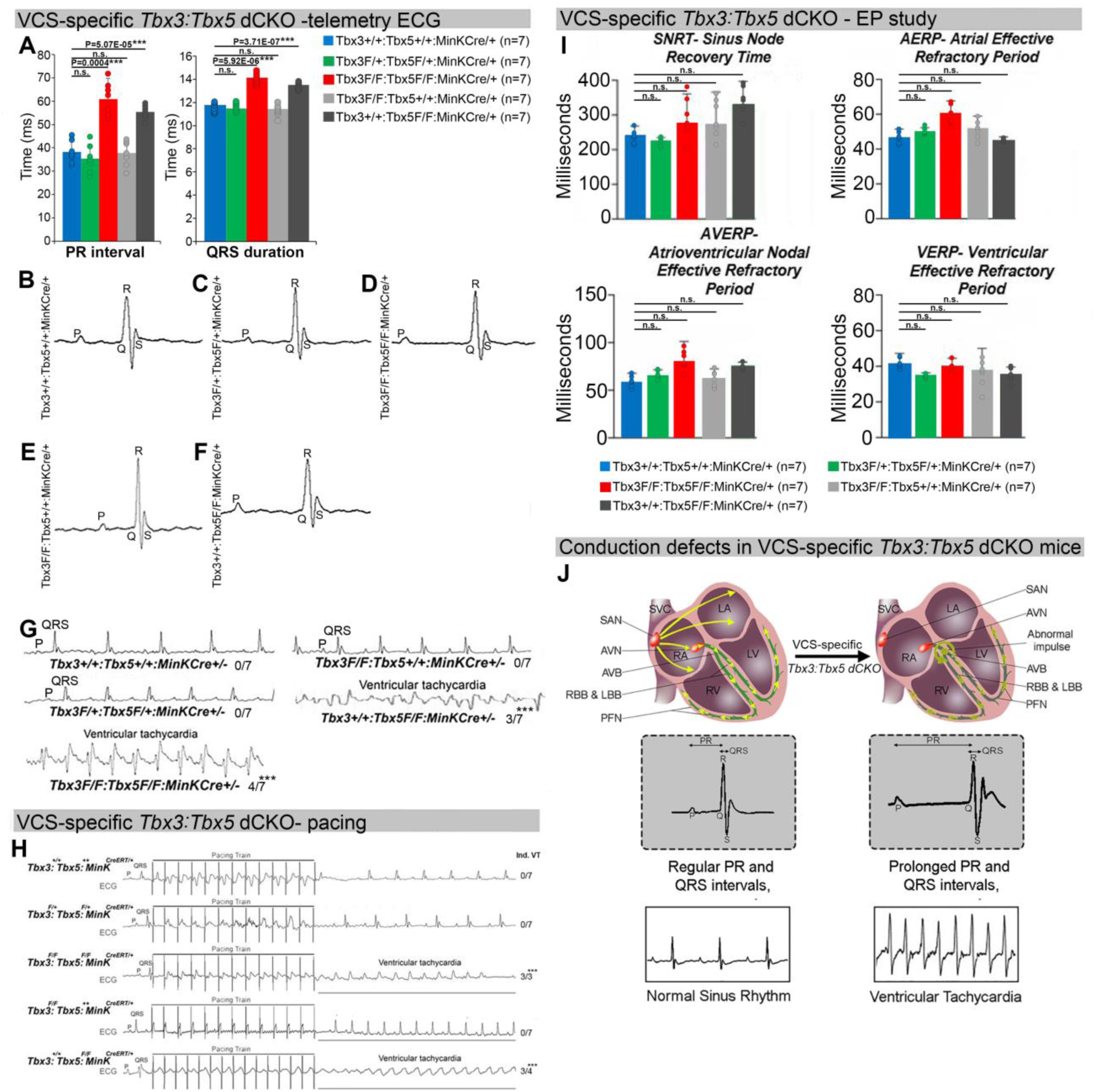
Arrhythmias and conduction abnormalities in mice with VCS-specific *Tbx3:Tbx5* double-conditional knockout. (A-F) VCS-specific *Tbx3:Tbx5* double- conditional knockout causes significant VCS conduction slowing in adult *Tbx3^fl/fl^;Tbx5^fl/fl^;R26^EYFP/+^;MinK^CreERT2/+^* mice. **(A)** PR **(left graph)** and QRS **(right graph)** intervals calculated from ambulatory telemetry ECG recordings in **(B-F)**. *Tbx3:Tbx5* double-conditional adult mice (*Tbx3^fl/fl^;Tbx5^fl/fl^;R26^EYFP/+^;MinK^CreERT2/+^*) displayed significant PR and QRS intervals prolongation compared to littermate controls (*Tbx3^+/+^;Tbx5^+/+^;R26^EYFP/+^;MinK^CreERT2/+^*) **(A left and right graphs, respectively)**. Data are presented as mean±SD. N=7 biological replicates/genotype, multiple testing correction using Benjamini & Hochberg procedure; *: Welch t-test P<0.05 & FDR<0.05; **: Welch t-test P<0.005 & FDR<0.05; ***: Welch t-test P<0.001 & FDR<0.05). **(B-F)** Representative ambulatory telemetry ECG of *Tbx3^+/+^;Tbx5^+/+^; R26^EYFP/+^;MinK^CreERT2/+^* **(B),** *_Tbx3_fl/+_;Tbx5_fl/+_;R26_EYFP/+_;MinK_CreERT2/+* **_(C),_** *_Tbx3_fl/fl_;Tbx5_fl/fl_; R26_EYFP/+_;MinK_CreERT2/+* **_(D),_** *_Tbx3_fl/fl_;Tbx5_+/+_;R26_EYFP/+_;MinK_CreERT2/+* **_(E),_** *_Tbx3_+/+_;Tbx5_fl/fl_;R26_EYFP/+_;MinK_CreERT2/+* **_(F)_** mice. **(G)** Simultaneous genetic removal of *Tbx3* and *Tbx5* from the adult VCS resulted in significantly increased episodes of spontaneous ventricular tachycardia. Episodes of spontaneous ventricular tachycardia were observed in 4 of 7 *Tbx3^fl/fl^;Tbx5^fl/fl^;R26^EYFP/+^;MinK^CreERT2/+^ mice versus* 0 of 7 littermate controls (*Tbx3^+/+^;Tbx5^+/+^;R26^EYFP/+^;MinK^CreERT2/+^*) in ambulatory studies. N=7 biological replicates/genotype; multiple testing correction using Benjamini & Hochberg procedure; * FDR of Welch t-test ≤0.05. **(H)** *Tbx3:Tbx5* double-conditional knockout mice (*Tbx3^fl/fl^;Tbx5^fl/fl^;R26^EYFP/+^;MinK^CreERT2/+^)* showed significantly increased susceptibility to ventricular tachycardia following burst stimulation in invasive electrophysiology studies (3 of 3 *Tbx3^fl/fl^;Tbx5^fl/fl^;MinK^CreERT2/+^* mice versus 0 of 7 control *Tbx3^+/+^;Tbx5^+/+^;R26^EYFP/+^;MinK^CreERT2/+^*.mice. Fisher’s exact test: *P<0.05; n=7 biological replicates/genotype). **(I)** Intracardiac electrophysiology detected no significant changes in SNRT, AERP, AVERP, and VERP recorded from experimental and control animals (n=7 biological replicates/genotype; multiple testing correction using Benjamini & Hochberg procedure; * FDR of Welch t-test ≤ 0.05). **(J)** Graphical summary of conduction defects observed in adult, VCS-specific *Tbx3:Tbx5*-deficient mice. Simultaneous genetic deletion of *Tbx3* and *Tbx5* from the mature VCS results in conduction slowing, prolonged PR and QRS intervals, as well as ventricular tachycardia. Abbreviations: AVB, atrioventricular bundle (also known as bundle of His); AVN, atrioventricular node; FDR, false discovery rate; LA, left atrium; LBB, left bundle branches; LV, left ventricle; PFN, Purkinje fiber network; RA, right atrium; RBB, right bundle branches; RV, right ventricle; SAN, sinoatrial node; SVC, superior vena cava, VCS, ventricular conduction system.

To distinguish a primary conduction system abnormality from a secondary conduction abnormality resulting from cardiac dysfunction or remodeling, we evaluated cardiac form and function at the time of arrhythmia assessment, 2-3 weeks post-tamoxifen treatment (Figure 3). Transthoracic echocardiography revealed no significant differences in left ventricular ejection fraction (LVEF) and fractional shortening (FS) between VCS-specific *Tbx3:Tbx5*-deficient (*Tbx3^fl/fl^;Tbx5^fl/fl^;R26^EYFP/+^;MinK^CreERT2/+^*) and control (*Tbx3^+/+^;Tbx5^+/+^;R26^EYFP/+^;MinK^CreERT2/+^*) mice *(LVEF: Tbx3:Tbx5* double-conditional knockout mice vs control mice P<0.05, n=7, Welch t test; and FS: *Tbx3:Tbx5* double-conditional knockout mice vs control mice P>0.05, n=7, Welch t test; Figure 3A, 3B and 3D). Histological examination of all four-chambers demonstrated no discernible differences between VCS-specific *Tbx3:Tbx5* double-knockout (*Tbx3^fl/fl^;Tbx5^fl/fl^;R26^EYFP/+^; MinK^CreERT2/+^*) and control (*Tbx3^+/+^;Tbx5^+/+^;R26^EYFP/+^; MinK^CreERT2/+^*) mice, nor between. the double-knockout (*Tbx3^fl/fl^;Tbx5^fl/fl^;R26^EYFP/+^; MinK^CreERT2/+^*) and single-knockout models for either *Tbx3* (*Tbx3^fl/fl^;Tbx5^+/+^;R26^EYFP/+^;MinK^CreERT2/+^*) or *Tbx5* (*Tbx3^+/+^;Tbx5^fl/fl^;R26^EYFP/+^;MinK^CreERT2/+^*). Ventricular muscle appeared normal without hypertrophy or myofibrillar disarray and no fibrosis was present (Figure 3G, 3I, 3J, and 3K, respectively). QRT-PCR analysis for fibrosis genes *Col1a1* (33–35) and *Postn* (35–38) further confirmed no fibrosis in VCS of *Tbx3:Tbx5*-deficient mice (Figure 3L). No contractile dysfunction, histological abnormalities, or increased expression of fibrosis genes were observed in VCS-specific *Tbx3:Tbx5* double-conditional heterozygous mice (*Tbx3^fl/+^;Tbx5^fl/+^;R26^EYFP/+^; MinK^CreERT2/+^*) (Figure 3A, 3C, 3H, and 3L). Taken together, these data indicate that the conduction defect and ventricular tachycardia observed in mice with VCS-specific *Tbx3:Tbx5* deletion (Figure 2) occur prior to the onset of left ventricular dysfunction or evidence of remodeling (Figure 3), implying a primary origin.

**Figure 3.**
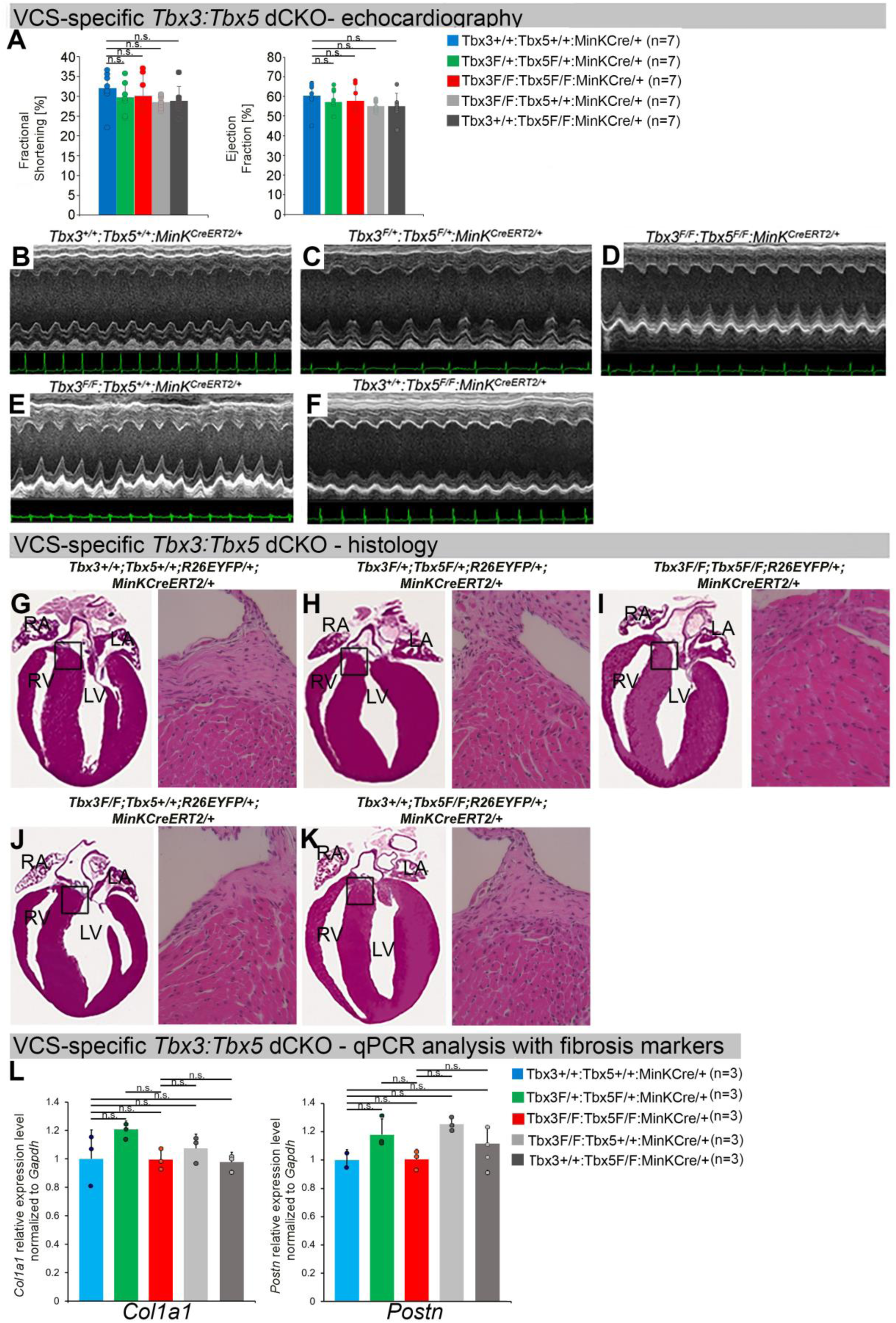
Cardiac function is preserved following double-conditional loss of *Tbx3* and *Tbx5* in the adult ventricular conduction system (VCS). (A) Left ventricular (LV) fractional shortening **(left graph)** and left ventricular (LV) ejection fraction **(right graph)** calculated from the M-mode ECGs in (B-F) revealed no contractile dysfunction in VCS- specific *Tbx3:Tbx5* double-conditional mutant mice (*Tbx3^fl/fl^;Tbx5^fl/fl^;R26^EYFP/+^; MinK^CreERT2/+^*). Data are presented as mean±SD. Welch *t* test: *P<0.05; ns, not significant; n=7 biological replicates/genotype. **(B-F)** Cardiac function, assessed by M-mode echocardiography from *Tbx3^+/+^;Tbx5^+/+^;R26^EYFP/+^;MinK^CreERT2/+^* (B), *Tbx3^fl/+^;Tbx5^fl/+^; _R26_EYFP/+_;MinK_CreERT2/+* _(C), *Tbx3*_*fl/fl_;Tbx5_fl/fl_;R26_EYFP/+_;MinK_CreERT2/+* _(D), *Tbx3*_*fl/fl_;Tbx5_+/+_;_ R26^EYFP/+^;MinK^CreERT2/+^* (E), *Tbx3^+/+^;Tbx5^fl/fl^;R26^EYFP/+^;MinK^CreERT2/+^* (F) mice shown above surface ECGs. No functional differences between mutant and control mice were detected. The most representative images for each genotype were utilized in the figure. n=7 biological replicates/genotype. **(G-K)** Histological examination of all four-chambers from *_Tbx3_+/+_;Tbx5_+/+_;R26_EYFP/+_;MinK_CreERT2/+* _(G), *Tbx3*_*fl/+_;Tbx5_fl/+_;R26_EYFP/+_;MinK_CreERT2/+* _(H), *Tbx3*_*fl/fl_;Tbx5_fl/fl_;R26_EYFP/+_;MinK_CreERT2/+* _(I), *Tbx3*_*fl/fl_;Tbx5_+/+_;R26_EYFP/+_;MinK_CreERT2/+* _(J),_ *Tbx3^+/+^;Tbx5^fl/fl^;R26^EYFP/+^;MinK^CreERT2/+^* (K) mice showed no histological abnormalities. The most representative images for each genotype were utilized in the figure. n=3-4 biological replicates/genotype. Boxed areas in **(G-K)** have been shown at higher magnification at their right sides. **(L)** qRT-PCR analysis for fibrosis genes *Col1a1* and *Postn* confirmed that there was no increase in expression of fibrosis markers in the VCS of *Tbx3:Tbx5*-deficient mice. Data are presented as mean±SD normalized to *Gapdh* and relative to *Tbx3^+/+^;Tbx5^+/+^;R26^EYFP/+^;MinK^CreERT2/+^* mice (set as 1). ns: FDR>0.05 in both Welch t-test and Wilcoxon test; n=2-3 biological replicates/genotype (VCS cardiomyocytes pooled from 30 mice per biological replicate). Histological examination original magnification: 2.5x, boxed area showed at the higher magnification: 40x. Abbreviations: FDR, false discovery rate; LA, left atrium; RA, right atrium; LV, left ventricle; RV, right ventricle.

To assess the hypothesis that *Tbx3* and *Tbx5* collectively promote VCS versus working myocardium phenotype, we conducted a transcriptional characterization of the adult VCS in *Tbx3^fl/fl^;Tbx5^fl/fl^;R26^EYFP/+^;MinK^CreERT2/+^* mutant mice compared to their *Tbx3^+/+^;Tbx5^+/+^;R26^EYFP/+^;MinK^CreERT2/+^* control littermates using three distinct sets of molecular markers by qRT-PCR (Figure 4A-C). The first set encompassed genes expressed throughout the entire conduction system (Pan-CCS) and implicated in slow-conducting nodal phenotype, such as *Hcn1, Hcn4, Cacna1d (Cav1.3), Cacna1g (Cav3.1d), Cacna1h (Cav3.2), Gjd3 (Cx30.2), and Gjc1 (Cx45)* (15, 30, 39–46) (Figure 4A). The second set included genes highly expressed in the fast-conducting VCS and important for VCS function, including *Gja5* (*Cx40), Scn5a (Nav1.5), Ryr2, Kcnk3 (Task- 1), Kcnj2 (Kir2.1), Kcnj3 (Kir3.1), Kcnj4* (IRK3), and *Kcnj12 (Kir2.2)* (Figure 4B) (39, 40, 46–50). The third set contained markers specifically present in the working myocardium but absent in the CCS, such as *Gja1* (*Cx43)* and *Smpx* (Figure 4C) (30, 39, 51, 52). VCS-specific *Tbx3:Tbx5*-deficient mice lost VCS expression profile of genes required for the fast ventricular conduction (Figure 4B) as well as genes normally expressed in whole CCS (Pan-CCS genes) (Figure 4A). In contrast, these mice obtained VCS expression of working myocardium-specific molecular markers (Figure 4C). Immunoblotting analysis confirmed transcriptional changes observed by qRT-PCR (Figure 4D). This molecular characterization indicated that the *Tbx3:Tbx5* double mutant VCS adopted a gene expression profile similar to wild-type working myocardial-like cells (Figure 4).

**Figure 4.**
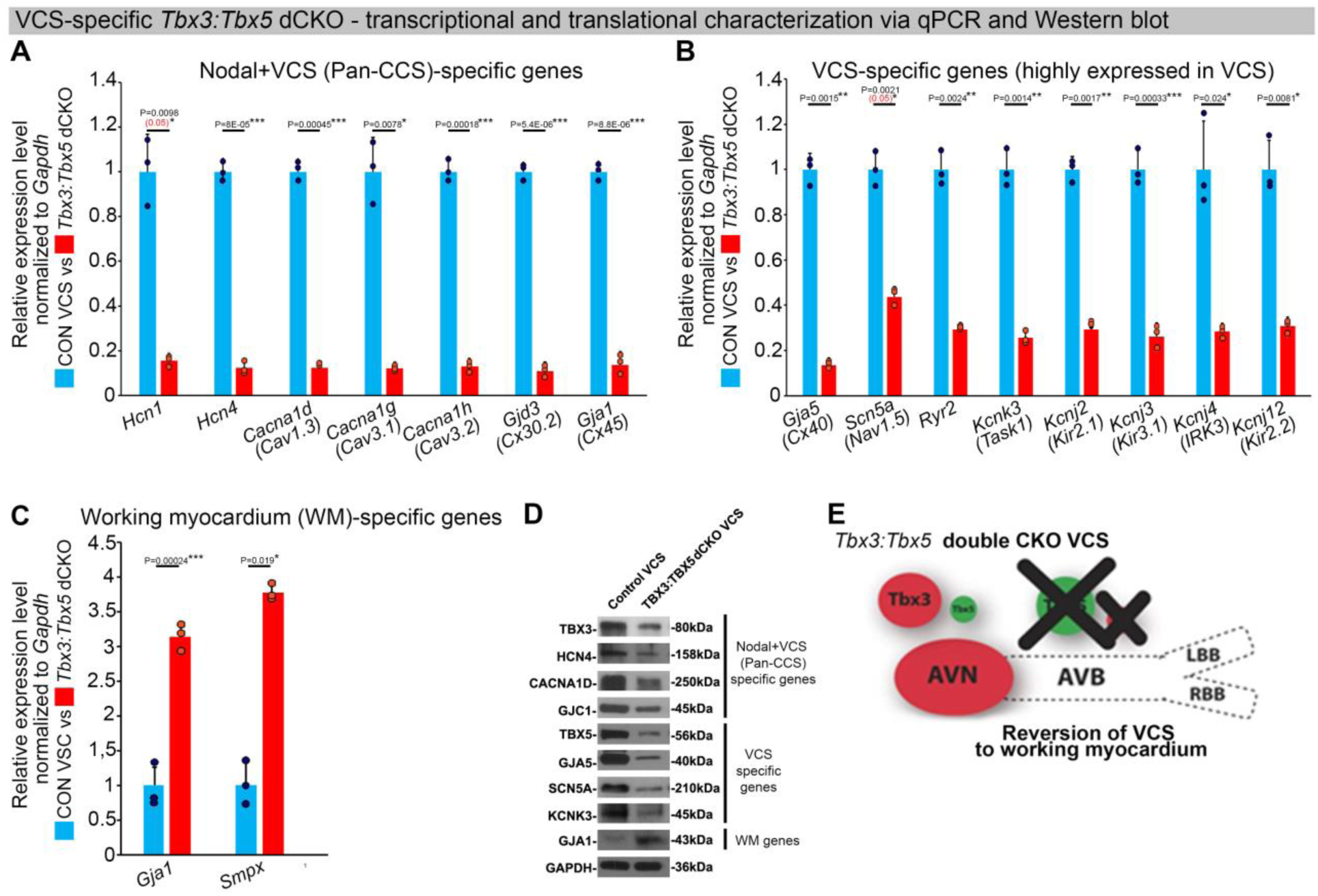
In the adult murine heart, *Tbx3* and *Tbx5* collectively **promote ventricular conduction system (VCS) versus working myocardium (WM) phenotype. (A-C)** qRT- PCR analysis of molecular changes driven by VCS-specific *Tbx3:Tbx5* double-conditional knockout in adult mice. Transcriptional characterization of the adult VCS in *Tbx3^fl/fl^;Tbx5^fl/fl^;R26^EYFP/+^;MinK^CreERT2/+^* mutant mice, compared to their *Tbx3^+/+^;Tbx5^+/+^;R26^EYFP/+^;MinK^CreERT2/+^* control littermates, was conducted using three distinct sets of molecular markers. **(A)** Genes expressed throughout the entire conduction system (Pan-CCS), implicated in the slow-conducting nodal phenotype. **(B)** Genes highly expressed in the fast-conducting VCS, critical for VCS function. **(C)** Markers specifically present in the working myocardium but absent in the CCS. VCS-specific *Tbx3:Tbx5*- deficient mice lost the VCS expression profile, including genes necessary for fast ventricular conduction **(B)** and those typically expressed in the entire CCS (Pan-CCS genes), essential for the slow conducting nodal phenotype **(A)**. In contrast, they acquired VCS expression of working myocardium-specific molecular markers important for working myocardial function **(C). (D)** Immunoblotting analysis confirmed transcriptional changes indicated by qRT-PCR analysis **(A-C)** in VCS-specific *Tbx3:Tbx5* double-conditional knockout in adult mice. **(E)** Graphical summary of transcriptional changes observed in VCS of VCS-specific *Tbx3:Tbx5*-deficient mice. Simultaneous genetic deletion of *Tbx3* and *Tbx5* from the mature VCS resulted in a transcriptional profile resembling that of ventricular working myocardium. QRT-PCR data are presented as mean±SD normalized to *Gapdh* and relative to *Tbx3^+/+^;Tbx5^+/+^;R26^EYFP/+^;MinK^CreERT2/+^* mice (set as 1). N=3 biological replicates/genotype (VCS cardiomyocytes pooled from 30 mice per biological replicate); multiple testing correction using Benjamini & Hochberg procedure. Significance was assessed by Welch *t* test (*FDR<0.05; **FDR<0.005; *** FDR<0.001) and confirmed by one-tail Wilcoxon test (P value in parentheses) when normally distribution was rejected. Abbreviations: AVB, atrioventricular bundle (also known as bundle of His); AVN, atrioventricular node; dCKO, double-conditional knock- out; FDR, false discovery rate; LBB, left bundle branches; RBB, right bundle branches; VCS, ventricular conduction system.

The impact of the *Tbx3:Tbx5* double-conditional knockout on electrical impulse propagation in the VCS of the heart was assessed with optical mapping of the anterior epicardial surface of the ventricles and right septal preparations where VCS function should be observed (Figure 5). Optical mapping records changes in transmembrane potential from multiple cells in tissue preparations, where ventricular septal optical action potentials (OAP) have two distinct action potential upstrokes. The first peak is a result of depolarization of the specialized fast VCS, followed by depolarization of ventricular working myocardium.

**Figure 5.**
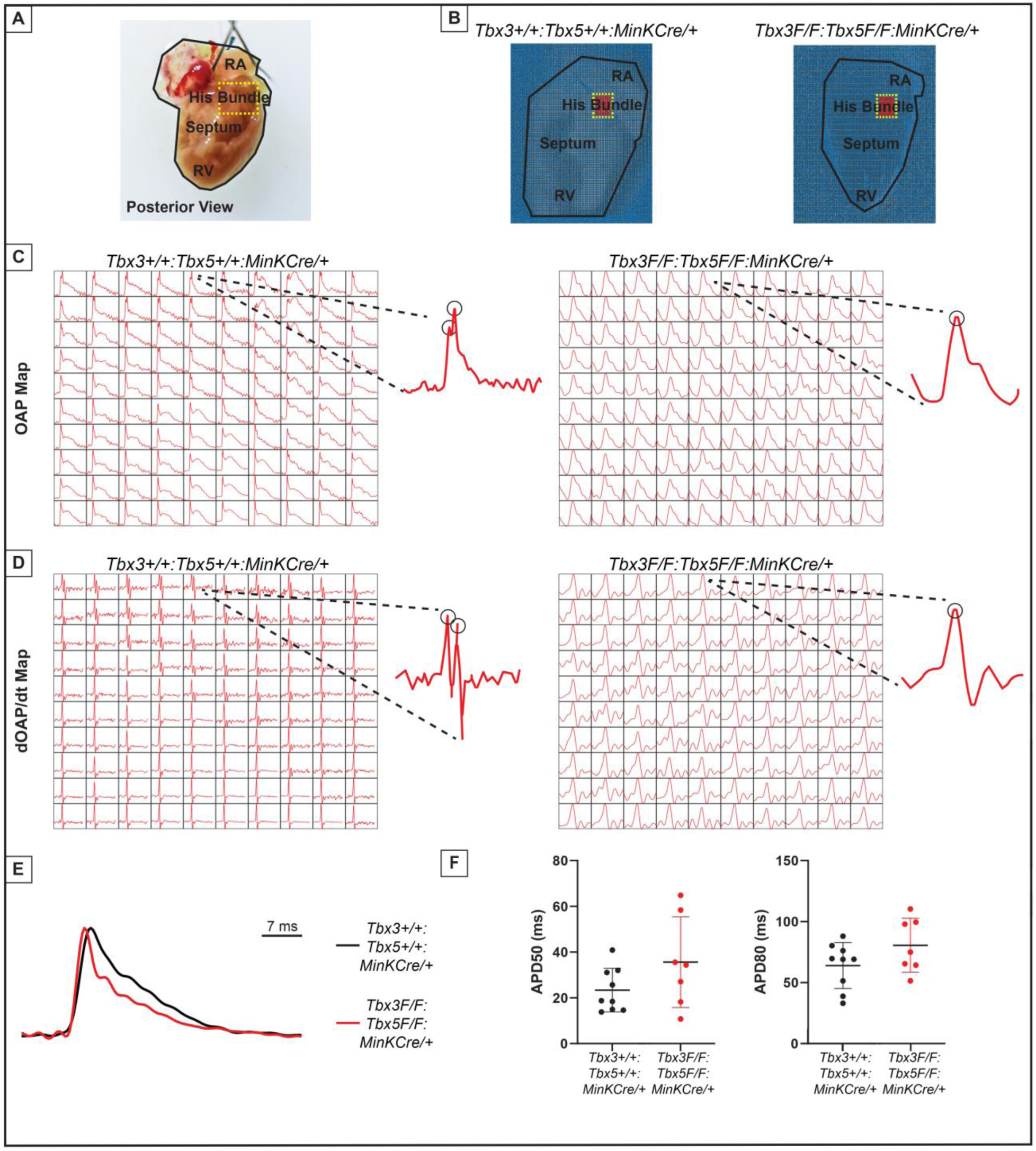
Loss of VCS optical action potential (OAP) morphology in *Tbx3:Tbx5* double-conditional knockout mice. **(A)** Schematic of the posterior view of a mouse heart with right ventricle (RV) free wall removed, highlight the RV, septum, His bundle, and right atria (RA). **(B)** Representative 100x100 OAP map of OAP recorded during sinus rhythm from *Tbx3:Tbx*5 double-conditional knockout mice and control littermates with the free wall removed. Rge region of the His bundle is highlighted in red. **(C)** Representative 10x10 OAP map from the region of the His bundle. **(D)** Representative 10x10 map of the first derivative of the OAP from the region of the His bundle. **(E)** Representative ventricular OAP from whole heart intact preparation from *Tbx3:Tbx*5 double-conditional knockout mice (red) and control littermates (black). **(F)** Quantification of APD50 and APD80 at a basic cycle length of 125 ms.

To visualize electrical impulse propagation in *Tbx3:Tbx5* double-conditional knockout mice, a 100x100 pixel data frame was plotted for the entire field of view of the right septal preparation (Figure 5A and B). For enhanced analysis of the VCS, the region encompassing the His bundle was distinguished in a red 10x10 area (Figure 5B). To specifically assess electrical impulse propagation within the His bundle, the same 10x10 pixel area representing this region was isolated from adult hearts of both control and *Tbx3:Tbx5* double- conditional knockout mice (Figure 5C). We observed only one OAP upstroke in *Tbx3:Tbx5* double-conditional knockout mice in contrast to control mice which showed two OAP upstrokes. The first derivative of the OAP (dOAP/dt) from the region of the His bundle was calculated and plotted in a dOAP/dt map further highlighting the number of temporally distinct upstrokes within the OAP (Figure 5D). In control mice, we observed two distinct depolarization upstrokes, one pertaining to the VCS and the second for working ventricular myocardium. However, in *Tbx3:Tbx5* double-conditional knockout mice only one maximum dOAP/dt was observed (Figure 5D). The OAP morphology observed in *Tbx3:Tbx5* double-conditional knockout mice suggested a loss or reduction of the specialized fast VCS conduction (Figure 5C and D).

OAPs from the anterior epicardial surface of the ventricles were compared between control littermates and *Tbx3:Tbx5* double-conditional knockout mice to assess changes in electrical impulse propagation in the ventricular working myocardium. Paced at a basic cycle length of 125 ms, control littermates and *Tbx3:Tbx5* double-conditional knockout mice did not exhibit significant differences in action potential duration at 50% (APD50) or 80% (APD80) repolarization (Figure 5E and 5F). These observations suggested that the electrical activity of the ventricular working myocardium was not appreciably altered in double knockout mice compared to controls. This finding is consistent with the lack of changes in the ventricular effective refractory period (VERP) observed in invasive electrophysiology (EP) studies of both control and *Tbx3:Tbx5* double-conditional knockout mice (Figure 3I).

We further predicted that the double knockout would not affect action potentials or conduction properties distal to the VCS (Figure 6). OAP and dOAP/dt maps were created to observe the effects on electrical impulse propagation distal to the His bundle on the right septal preparation (Figure 6), similar to the methods applied to create OAP and dOAP/dt maps in the region of the His bundle (Figure 5), with the key difference being that the signals were recorded from the red 10x10 pixel regions plotted in the working ventricular myocardium distal to the His bundle instead of in the area of the His bundle (Figure 6A and B versus Figure 5A and B, respectively). In both control littermates and *Tbx3:Tbx5* double-conditional knockout mice, only one action potential upstroke and one dOAP/dt maximum are observed which indicates that most of this region consists of the ventricular working myocardium (Figure 6C and D, respectively). In summary, the significant remodeling of electrical impulse propagation due to *Tbx3:Tbx5* double-conditional knockout was observed by the loss of a distinct fast VCS impulse in the region of the His bundle, but not on the anterior epicardial surface of the ventricles or distal to the His bundle on the right ventricular septum (Figure 5 and 6).

**Figure 6.**
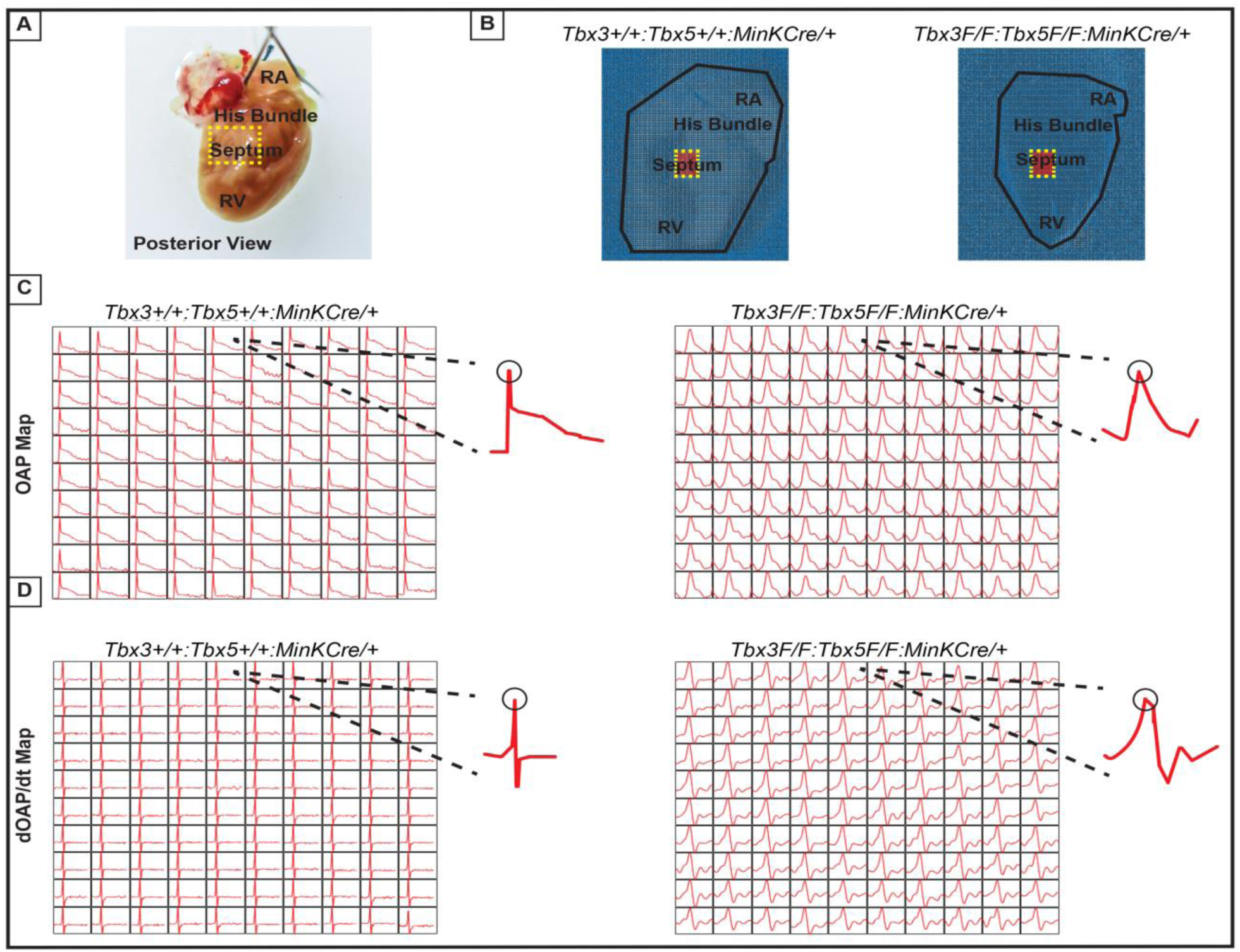
Ventricular optical action potentials (OAPs) distal from His bundle have only 1 OAP upstroke. **(A)** Schematic of the posterior view of mouse heart with right ventricle (RV) free wall removed. **(B)** Representative 100x100 pixel OAP map recorded during sinus rhythm from *Tbx3:Tbx*5 double-conditional knockout mice and control littermates with RV free wall removed. The region of the working ventricular myocardium distal from the His bundle is highlighted in red. **(C)** Representative 10x10 pixel OAP map from the region distal to the His bundle. **(D)** Representative 10x10 dOAP/dt map from the region distal to the His bundle.

## Discussion

Our study investigated the impact of compound *Tbx3:Tbx5*-deficiency on mature VCS function and molecular identity. Using a double-conditional knockout strategy, both genes were targeted specifically in the adult VCS. Loss of *Tbx3* and *Tbx5* expression in the mature VCS led to profound conduction defects, characterized by prolonged PR interval and QRS duration, increased susceptibility to ventricular tachycardia, and sudden death. These alterations were observed in the absence of discernible changes in cardiac contractility or histological morphology, indicating a primary conduction system defect. Molecular characterization of the adult VCS unveiled an altered gene expression profile in *Tbx3:Tbx5* double-conditional knockout mice, suggesting a transition from the distinctive fast VCS transcriptional profile to that resembling ventricular working myocardium. Optical mapping demonstrated loss of the specialized fast VCS function in *Tbx3:Tbx5* double-conditional knockout mice, further suggesting that this region acquired an electrophysiological phenotype similar to the ventricular working myocardium.

Our previous research (1, 3, 11, 12) and published literature (7, 15, 20, 26, 53, 54) have suggested that the balance between *Tbx3* and T*bx5* expression determines the regional specialization of the mature central cardiac conduction system (Figure 7). *Tbx3* expression dominates in nodal myocardium, imparting nodal physiologic characteristics. *Tbx5* expression dominates in fast VCS myocardium, where a T-box-dependent fast conduction system network drives physiologically dominant fast conduction physiology, overriding nodal physiology (11) (Figure 7). This model accurately predicts the outcome of targeted manipulations, such as adult VCS-specific removal of TBX5 or overexpression of TBX3, which convert the fast VCS into a slow nodal-like system (11) (Figure 7).

**Figure 7.**
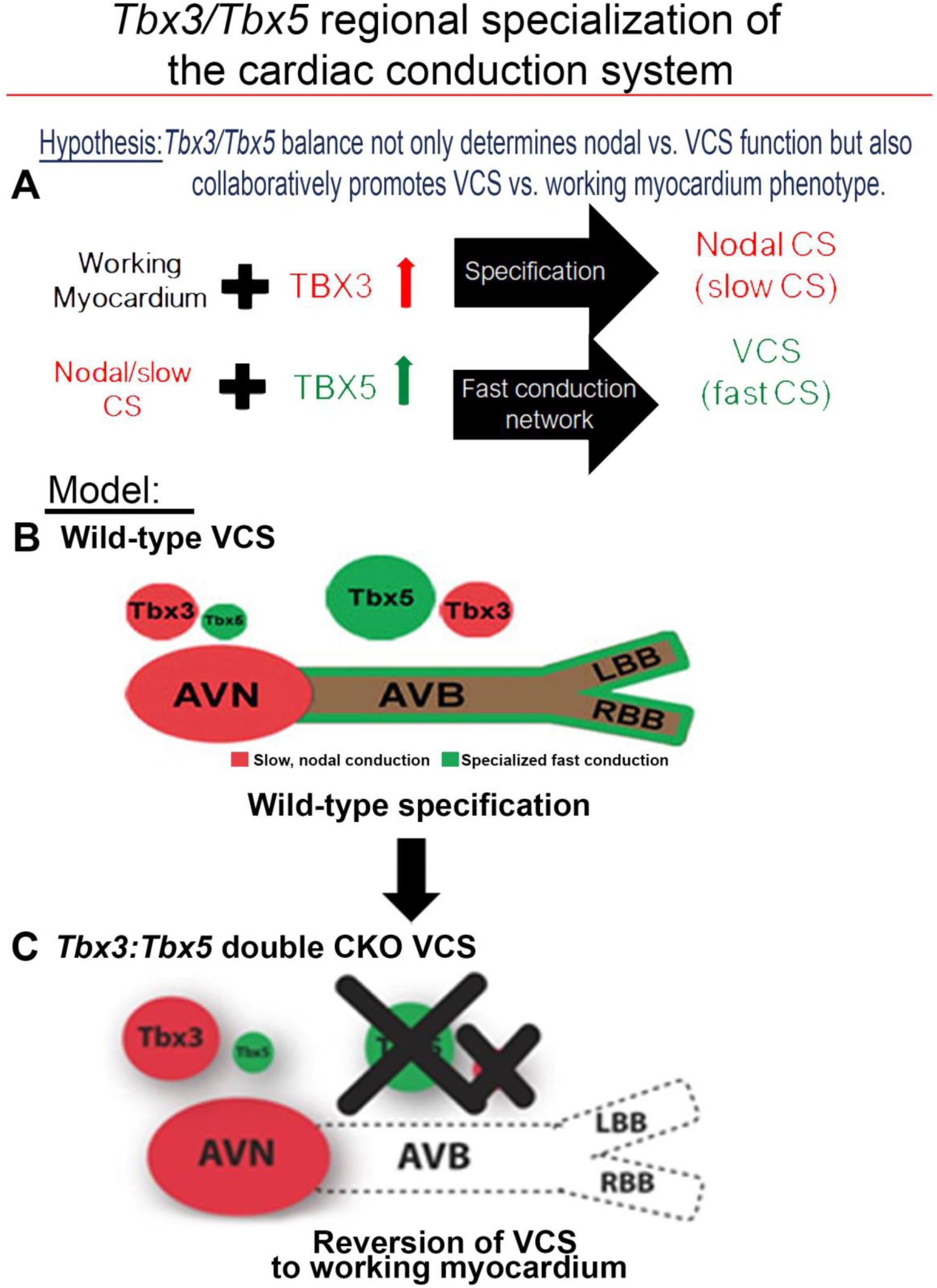
*Tbx3* and *Tbx5* play distinct roles in the adult VCS while collectively promoting CCS regional identity - a model elucidating our hypothesis for *Tbx3/Tbx5* dose-dependent CCS regional specialization. (A) The *Tbx3/Tbx5* balance not only governs nodal versus ventricular conduction system (VCS) function but also collaboratively promotes the VCS versus working myocardium (WM) phenotype. Specifically, a high level of *Tbx3* is linked to the specialization to the nodal conduction system, while an elevated *Tbx5* level in nodal cells activates local expression of the *Tbx5*- dependent fast conduction network, resulting in the generation of VCS. **(B)** CCS regional specialization is driven by local expression of *Tbx5*-dependent fast conduction network in the VCS, which overlaps underlying Pan-CCS expression of nodal, slow conduction network. **(C)** VCS-specific simultaneous genetic removal of both the *Tbx3* and *Tbx5* transcription factors transforms the fast-conducting, adult VCS into cells resembling working myocardium, thereby shifting them from conduction to non-conduction myocytes. Therefore, within the adult CCS, the *Tbx3* and *Tbx5* expression levels are crucial not only for normal fast versus slow conduction system identity but also for maintaining the conduction versus contraction specialization of the VCS. AVB indicates atrioventricular bundle; AVN, atrioventricular node; CCS, cardiac conduction system; CKO, conditional knock-out; LBB, left bundle brunch; RBB, right bundle brunch; VCS, ventricular conduction system.

The adult CCS organization model posits that *Tbx3* and *Tbx5* coordinately establish CCS characteristics and suggests their compound necessity to maintain specialized VCS identity (Figure 7), a hypothesis that remained untested. This model predicted that VCS-specific genetic ablation of both the TBX3 and TBX5 transcription factors would transform fast-conducting adult VCS into cells resembling working myocardium, eliminating specialized CCS identity (Figure 7). Testing this model necessitated the generation of a *Tbx3:Tbx5* double-conditional knockout allele due to their proximal chromosomal location in *cis* on mouse Chr5. VCS-specific double knockout mice showed a notable deceleration in VCS conduction, manifested by prolonged PR and QRS intervals, along with increased susceptibility to ventricular tachycardia (VT). A similar functional phenotype was observed in single deletion of *Tbx5* from the adult VCS, leading to the transformation of the fast VCS into a nodal-like phenotype (1, 11). However, we predicted that the nodal-like characteristics of the *Tbx5*-ablated VCS were due to retained expression of *Tbx3.* In fact, the autonomous beating and impulse initiation observed in the *Tbx5*-mutant VCS (1, 11) were absent from the double *Tbx3:Tbx5* mutant VCS, further suggesting the transformation from a nodal-like to an inert myocardial functionally and a shift from conduction towards simple working myocardium.

Molecular studies support a transformation from conduction to working myocardium in the *Tbx3:Tbx5* double VCS knockout. Previous studies investigating the roles of *Tbx3* and *Tbx5* in maintaining adult VCS identity demonstrated that *Tbx3* deletion resulted in the silencing of Pan-CCS gene expression in the atrioventricular conduction system (7, 15, 20, 26, 53). Alternately, specific deletion of *Tbx5* from the adult VCS led to the decreased expression of VCS-specific markers while Pan-CCS markers remained unchanged, indicating a transformation towards a nodal-like transcriptional phenotype in the absence of *Tbx5* (1, 11). Consistently, a similar transformation was induced by *Tbx3* overexpression in the adult VCS. These findings underscored the significance of the *Tbx3:Tbx5* ratio in preserving the molecular and functional characteristics of the fast versus slow conduction system. In *Tbx3:Tbx5* double VCS knockout, we observed a reduction in the expression of both fast VCS markers and Pan-CCS markers transcribed throughout the entire CCS. As expected, not all VCS markers were ablated in VCS-specific *Tbx3:Tbx5* mutants. A significant portion of VCS fast conduction markers are also transcribed in ventricular working myocardium, albeit at lower levels than in the VCS, and are crucial for normal working myocardial function, e.g., *Ryr2, Scn5a,* and *Kcnj2* (39, 46, 49). Consistent with a shift from fast VCS to working myocardium, their expression levels are present, but at significantly lower levels in the double knockout mutants than in normal VCS. Furthermore, the expression of markers specific for working myocardium, which are normally excluded in the VCS, emerged in the VCS of *Tbx3:Tbx5* double mutant. These observations are consistent with a transcriptional shift from VCS conduction cardiomyocytes to working myocardium-like characteristics in the absence of both *Tbx3* and *Tbx5*.

Our study highlights the critical role of *Tbx3* and *Tbx5* in maintaining the specialized electrical properties of the VCS. Comparative analysis of the phenotypes observed in single *Tbx3* or *Tbx5* knockouts and the *Tbx3:Tbx5* double-conditional knockout supports a coordinated function for these transcription factors in preserving VCS functionality. In a normal mammalian heart, electrical or optical recordings from the proximal part of the ventricular conduction system typically display two distinct electrical excitations, reflected as two spikes in electrograms or two upstrokes in optical action potentials. These signals correspond to the sequential activation of the VCS, followed by ventricular excitation, a phenomenon well documented in both basic and clinical studies across species, including mice and humans. Previous whole-heart electrophysiology and optical mapping studies demonstrated that the single conditional knockout of *Tbx5* in the adult VCS caused significant ventricular conduction slowing (1), accompanied by a phenotypic shift of VCS cells toward a pacemaker-like cell, evidenced by ectopic beats originating in the ventricles, retrograde activation, and inappropriate automaticity (11). Whole-cell patch clamp recordings of *Tbx5*-deficient VCS cells further supported these findings, revealing action potential characteristics resembling pacemaker cells, including a slower upstroke (phase 0), prolonged plateau (phase 2), delayed repolarization (phase 3), and enhanced phase 4 depolarization (11). In contrast, reduced expression of *Tbx3* in the *Tbx3* haploinsufficiency model resulted in AV bundle hypoplasia, PR interval shortening, and prolonged QRS duration (53). Optical mapping in these mice showed delayed apical activation with multiple small breakthroughs separated by regions of delayed activation. This fragmented activation pattern suggests that the reduced *Tbx3* expression impaired the homogeneity and efficiency of ventricular electrical impulse propagation due to AV bundle and bundle branch hypoplasia (53). In the *Tbx3:Tbx5* double-conditional knockout mice, we observed a complete loss of fast VCS conduction, similar to the phenotype of the *Tbx5* single conditional knockouts. However, unlike in the *Tbx5* single CKO, the VCS in *Tbx3:Tbx5* double mutants did not acquire pacemaker-like activity. Instead, optical mapping revealed a single upstroke in the His bundle region of *Tbx3:Tbx5* double-conditional knockout mice, indicating the absence of rapid AV bundle electrical impulse propagation. Functionally, this transformation resulted in significantly slowed ventricular activation, without the retrograde activation observed in the *Tbx5* single conditional knockouts or the rapid electrical impulse propagation typically associated with an intact VCS. The homogenous morphological and molecular transformation of His bundle and VCS cells in the *Tbx3:Tbx5* double-conditional knockout mice rendered the electrical properties of the VCS-located cells functionally indistinguishable from those of ventricular working myocardium. Our data including gene expression analysis of the double mutant VCS are most consistent with the transition of double mutant VCS cells toward a working myocardium phenotype.

These findings emphasize the interdependent roles of *Tbx3* and *Tbx5* in maintaining the specialized electrical properties and identity of the VCS, distinguishing it from non-specialized ventricular working myocardium (Figure 7). While *Tbx5* alone is critical for suppressing pacemaker-like characteristics and preserving fast conduction properties, *Tbx3* plays a pivotal role in ensuring the structural and functional integrity of the AV and BB conduction pathways. The simultaneous deletion of both *Tbx3* and *Tbx5* disrupts these regulatory mechanisms, leading to a loss of VCS identity and a functional shift toward a working myocardium phenotype. Our study combined with prior literature (1, 7, 11, 15, 26, 53, 54) indicates that the concurrent presence of both *Tbx3* and *Tbx5* is necessary for maintaining VCS identity in the adult heart, ensuring efficient electrical conduction and preventing the transition of VCS cells into a working myocardium-like state (Figure 7).

## Methods

### Data Availability

The authors declare that all supporting data and materials presented within this manuscript and its Online Supplemental Materials are available from the corresponding author upon reasonable request.

### Experimental animals

All animal experiments were performed under the University of Chicago Institutional Animal Care and Use Committee (IACUC) approved protocol (ACUP no. 71737) and in compliance with the USA Public Health Service Policy on Humane Care and Use of Laboratory Animals. *MinK^CreERT2^* [*Tg(RP23-276I20-MinKCreERT2)*] and *Tbx3^fl/fl^* mice have been reported previously (8, 26).

*Tbx3:Tbx5* double floxed mouse line was generated by University of Utah Core Research Facility using CRISPR/Cas9. Guide RNA (sgRNA) constructs were designed with software tools (ZiFiT Targeter (55) and crispr.genome-engineering.org) predicting unique target sites throughout the mouse genome (Supplementary Figure 3 - 5). The sgRNA constructs were transcribed *in vitro* using MEGAshortscript T7 (Invitrogen AM1354) and mMessage Machine T7 transcription kit (Invitrogen AM1344) according to manufacturer instructions. The strategy to generate mouse founders involved a single-step microinjection into one-cell *Tbx3* floxed zygotes (on a mixed background) with 10 ng/µL of each sgRNA (TBX5-I2-S22 and TBX5-I3-S31, Supplementary Figure 3 - 5), 30 ng/µL of Cas9 protein, and 10 ng/µL of a long ssDNA donor. The donor contained two lox2272 sites in cis, spanning partial intron 2, entire exon 3, and partial intron 3 of the mouse *Tbx5* gene, along with 100/150 bp 5’/3’ homology arms (Supplementary Figure 4).Founders were validated by PCR, restriction enzyme digestion, and Sanger sequencing (Supplementary Figure 1B, C, and D). Founders were backcrossed with wild-type *CD1 IGS* mice (Charles River Lab, USA) to confirm germline transmission of the *CRISPR/Cas9*-generated compound *Tbx3:Tbx5* double-floxed allele and obtain the F1 generation. F1 mice were then interbred to establish a stable *Tbx3:Tbx5* double-floxed mouse line. Downstream experiments were performed on F4-F6 mice.

To simultaneously conditionally delate the *Tbx3* and *Tbx5* genes specifically from the adult VCS, we crossed our *Tbx3:Tbx5* double-floxed mouse line (*Tbx3^fl/fl^;Tbx5^fl/fl^*) with a VCS-specific tamoxifen (TM) inducible *Cre* transgenic mouse line (*MinK^CreERT2^* [*Tg(RP23-276I20-MinKCreERT2*; (8)) (Figure 1A). All mice were maintained on a mixed genetic background. Tamoxifen (MP Biomedical) was administered at a dose of 0.167 mg/g body weight for 5 consecutive days by oral gavage at 6 weeks of age and then mice were evaluated at 9 weeks of age, as previously described (1, 8, 11). Age-, gender-, and genetic strain-matched controls were used in all experiments to account for any variations in data sets across experiments. Mice were bred and housed in specific pathogen–free conditions in a 12-hour light/12-hour dark cycle and allowed ad libitum access to standard mouse chow and water. Mice requiring medical attention were provided with appropriate veterinary care by a licensed veterinarian and were excluded from the experiments described. No other exclusion criteria were applied.

All experiments and subsequent analysis were conducted in a blinded fashion, with animals randomly assigned to experimental groups. Following genotyping, mice were randomly allocated to the studies based on their genotypes. Subsequently, their identities were anonymized using a numerical code to ensure that all experiments and analyses were performed in a blinded manner. Both, male and female animals have been used in our studies in the ratio of 41%/59%, based on availability of relatively rare compound genotypes, respectively.

### Echocardiography studies

Transthoracic echocardiography in mice was conducted under inhaled isoflurane anesthesia administered through a nose cone. Prior to imaging, chest hairs were removed using a topical depilatory agent. Limb leads were affixed for electrocardiogram gating, and animals were imaged in the left lateral decubitus position with a VisualSonics Vevo 770 machine using a 30-MHz high-frequency transducer. To ensure stability, body temperature was carefully maintained using a heated imaging platform and warming lamps. Two-dimensional images were meticulously recorded in parasternal long- and short-axis projections, accompanied by guided M-mode recordings at the midventricular level in both views. Left ventricular (LV) cavity size and percent fractional shortening were measured in at least 3 beats from each projection and averaged. M-mode measurements were employed to ascertain LV chamber dimensions and percent LV fractional shortening, calculated as ([LVIDd – LVIDs]/LVIDd), where LVIDd and LVIDs represent LV internal diameter in diastole and systole, respectively.

### Surface electrocardiography (ECG)

9 weeks old, tamoxifen treated control and mutant mice were anesthetized with a mixture of 2-3% isoflurane in 100% oxygen. Anesthetized mice were secured in a supine position on a regulated heat pad while lead I and lead II ECGs were recorded using platinum subdermal needle electrodes in a 3-limb configuration. Core temperature was continuously monitored using a rectal probe and maintained at 36–37 °C throughout the procedure. ECG data were collected and analyzed using Ponemah Physiology Platform (DSI) software and an ACQ-7700 acquisition interface unit (Gould Instruments, Valley View, OH, USA). Key parameters derived from the ECG measurements included: Heart rate (HR), PR interval (from the beginning of the P wave to the beginning of the QRS complex), and QRS complex duration.

### Telemetry ECG analysis

9 weeks old, tamoxifen treated control and mutant mice were anesthetized with 2-3% isoflurane in 100% oxygen, and wireless telemetry transmitters (ETA-F10; DSI) were surgically implanted in the back with leads tunneled to the right upper and left lower thorax, as previously described (1, 56). Following 24-hour recovery period after surgical instrumentation, heart rate and PR and QRS intervals were calculated using Ponemah Physiology Platform (DSI) from 48-hour recordings.

### Catheter-based intracardiac electrophysiology

Detailed protocols for invasive electrophysiology studies (EP) have been previously described (57, 58). Briefly, 9 weeks old, tamoxifen treated control and mutant mice were anesthetized using 2-3%isoflurane in 100% oxygen. Then, a 1.1-Fr octapolar electrode catheter (EPR-800; Millar Instruments) was advanced via a right jugular venous cut-down to record right atrial (RA), His bundle, and right ventricular (RV) potentials, as well as to perform programmed electrical stimulation. Signals were identified through alignment with simultaneous surface electrocardiography (ECG) using subcutaneous needle electrodes in a Lead II configuration. “Near-Field” and “Far-Field” signals were identified based on ECG alignment, signal deflection upstroke speed and total signal duration. Standard tachycardia induction protocols included an 8-beat drive train with beats 80-120ms apart (S1), followed by 5 beats (S2) at 50ms apart (penta-extrastimulus, PES). Two attempts at this PES protocol were carried out. Mice also underwent single extrastimulus testing (SES) with 8 beats 80-120ms apart (S1) followed by a single S2 at 50ms. This SES protocol was carried out 5 separate times per mouse. If these two protocols in the right atrium and right ventricle (separately) failed to initiate their respective tachycardias, the study was deemed negative. S1 drive train intervals varied slightly due to the presence of the AV Wenckebach block at faster pacing rates in some mice; the S1 interval was lengthened to prevent this during the drive train.

Additional atrial and ventricular pacing protocols were carried out to obtain atrial, atrio-ventricular, and ventricular effective refractory periods (AERP, AVERP, VERP) as well as sinus node recovery time, as described previously (57–60). Effective refractory periods were measured using 8-beat S1 drive trains of 100 ms followed by single extra- stimulus.

### ECG and Optical mapping

#### Electrocardiogram (ECG) Acquisition and Analysis

ECGs were recorded in conscious *Tbx3:Tbx5* double-conditional knockout mice and control littermates using the ecgTUNNEL device (emka Technologies) 2 weeks after tamoxifen treatment. Mice were positioned in the tunnel and ECGs were recorded for 5 minutes at a sampling rate of 1 kHz using lead I. ECGs were analyzed using a custom MATLAB program to measure P, and QRS durations, as well as PR, QT and RR intervals (61, 62) (Supplementary Figure 6).

#### Langendorff Perfusion

*Tbx3:Tbx5* double-conditional knockout mice (n=8) and control littermates (n=10) were deeply anesthetized with isoflurane. The heart was quickly excised following cervical dislocation and thoracotomy. The aorta was cannulated, and the heart was retrogradely perfused with warmed (37°C) and oxygenated (95% O2 and 5% CO2) modified Tyrode’s solution (in mM, NaCl 130, NaHCO3 24, NaH2PO4 1.2, MgCl2 1, Glucose 5.6, KCl 4, and CaCl2 1.8) at a pH of 7.4. The heart was placed in a constant-flow (1.0 - 2.0 ml/min) Langendorff Perfusion system, laying horizontally in a tissue bath.

#### Ventricular Optical Mapping

The Langendorff Perfused heart was electromechanically uncoupled by 15 μM of blebbistatin (Cayman Chemicals 13186) perfusion. A voltage-sensitive fluorescent dye, 80 µM di-4-ANEPPs (ThermoFisher Scientific D1199), was administered through the dye injection port. The heart was illuminated using a 520 ± 5 nm (Prizmatix, UHP-Mic-LED- 520) wavelength light source to excite di-4-ANEPPs. Emitted photons were captured using complementary metal-oxide semiconductor (CMOS) cameras (MiCAM, SciMedia). The stimulation threshold, where the heart would capture the stimuli and action potential 1:1, was determined using a point source platinum electrode placed on the anterior surface of the heart on the anterior epicardial surface of the ventricles. Pacing was applied at 1.5x the threshold amplitude to maintain 1:1 capture over the duration of the experiment. Optical recordings were analyzed using Rhythm 1.2 to analyze transmembrane potential (63). Action potential duration (APD) at 50% and 80% repolarization were calculated for the ventricles of the intact whole heart preparation.

#### Right Septal Preparation Optical Mapping Experiments

Following intact whole heart *ex vivo* optical mapping, the heart was removed from the tissue bath for the dissection of the right ventricular (RV) free wall to expose the RV septal surface (64). The heart was quickly returned to the tissue bath and the RV septal surface was focused into the field of view. Sinus rhythm optical recordings were acquired.

A custom MATLAB program was written to plot individual pixels in a 100x100-pixel image stack. A Butterworth filter of order 5 was applied to provide temporal filtering (65). The location of the His bundle was identified as the region at the base of the interventricular septum (a red 10x10 pixel area in Figure 5B). Signals from the working ventricular myocardium were recorded from the 10x10 pixel region plotted in the working ventricular myocardium distal to the His bundle (Figure 6B).

### Isolation of adult VCS cardiomyocytes and cell sorting

Adult mouse EYFP-positive VCS cardiomyocytes were isolated using the method described by Mitra and Morad (66). Briefly, 9 weeks old tamoxifen treated control and *Tbx3:Tbx5* double-conditional mutant mice were heparinized (100 units IP) and anesthetized with pentobarbital (50 mg/kg of body weight). Hearts were excised and mounted on a Langendorff apparatus, then perfused with Ca2+-free Tyrode’s solution with collagenase B and D (Roche Chemical Co.) plus protease (Fraction IV, Sigma Chemical Co.). When the hearts appeared pale and flaccid, they were removed from the Langendorff apparatus and a tip of ventricular septum below the AV annulus was microdissected out (1, 2, 67) and kept in Ca2+-free Tyrode’s solution with 1 mg/ml of bovine serum albumin (Fraction XIV, Sigma Chemical Co.). The intraventricular sections were then minced into small piece ∼1 x 1 x 1 mm and then gently triturated with a Pasteur pipette to dissociate individual VCS myocytes. Propidium iodide (ThermoFisher Scientific) was added immediately before FACS to facilitate live/dead discrimination. Cells were sorted on a FacsAria flow cytometer (BD Biosciences) located at the University of Chicago Flow Cytometery Core using Influx software. Samples from wild-type age-matched hearts were used for gating. Samples were gated to exclude debris and cell clumps. Fluorescent cells were collected into ice-cold RNase-free PBS and processed for RNA extraction and protein isolation.

### RNA isolation and QRT-PCR

Total RNA was isolated from EYFP-positive VCS cardiomyocytes sorted from 9 weeks old control and VCS-specific *Tbx3:Tbx5*-deficient mice, which had received tamoxifen at 6 weeks of age. RNA was also extracted from atrial and ventricular tissues dissected from the same mice. All RNA extractions were performed using RNeasy Mini Kit (Qiagen), followed by DNase treatment according to the manufacturer’s instructions.

Reverse transcription reaction was carried out using the SuperScript III First-Strand Synthesis SuperMix for quantitative RT-PCR (Invitrogen) as per the manufacturer’s recommendations. QRT-PCR was performed using the POWER SYBR Green PCR master mix from Applied Biosystems and run on an Applied Biosystems AB7500 machine in 96 well plates. The relative gene expression level was calculated by the ΔΔCt method (68) using glyceraldehyde-3-phosphate dehydrogenase (Gapdh) gene expression level as internal control. The data presented are the average of three independent experiments.

#### Protein isolation and Western blotting

Total protein was isolated from EYFP-positive VCS cardiomyocytes sorted from 9 weeks old control and VCS-specific *Tbx3:Tbx5*-deficient mice that had been administered tamoxifen at 6 weeks of age. Protein was also obtained from atrial and ventricular tissues dissected from the same animals. The tissues were snap-frozen in liquid nitrogen, pulverized, and homogenized in RIPA buffer (50 mM Tris-HCl pH 8, 150 mM NaCl, 1% Triton-X, 0.5% sodium deoxycholate, 0.1% SDS, 5 mM EDTA) with 1 Roche EDTA-Free complete protease inhibitor tablet per 50 mL of buffer. Samples were tumbled for 1 hour at 4°C and then centrifuged for 10 minutes at 13,200 x g. Protein concentration was determined using the BCA assay (Pierce) with BSA as a standard.

For Western blot analysis, 25 µg of protein was diluted in Laemmli buffer, heated at 70°C for 10 minutes, and subjected to SDS-PAGE on 4-20% TGX gels (Bio-Rad). Proteins were then transferred to nitrocellulose membranes, blocked with 5% milk in TBS-T, and incubated overnight at 4°C with primary antibodies diluted in 2.5% milk in TBS-T. The primary antibodies used were: goat polyclonal anti-TBX3 (Santa Cruz Biotechnology, sc- 31656, 1:250), rabbit polyclonal anti-HCN4 (Millipore, AB5808, 1:500), rabbit polyclonal anti-CAV1.3/CACNA1D (Alomone, ACC-005, 1:200), rabbit polyclonal anti-Cx45/GJC1 (Thermo Fisher, PA5-77357, 1:250), sheep polyclonal anti-TBX5 (R&D, AF5918, 1:200), rabbit polyclonal anti-CX40/GJA5 (Zymed/Invitrogen, 36-4900, 1:500), rabbit polyclonal anti-NAV1.5/SCN5A (Alomone, ASC-005, 1:200), mouse monoclonal anti- KCNK3/TASK1 (Abcam, ab186352, 1:1000), rabbit polyclonal anti-CX43/GJA1 (Cell Signaling Technology, 3512, 1:1000), and mouse monoclonal anti-GAPDH (Abcam, ab8245, 1:1000). After rinsing in TBS-T, membranes were incubated for one hour at RT with secondary antibodies diluted in 2.5% milk in TBS-T, rinsed again, and visualized using enhanced chemiluminescence reagents (Pierce ECL/ECL Plus, Thermo Fisher Scientific) and Kodak X-OMAT film. Results were normalized to GAPDH loading control and then quantified using ImageJ software (69, 70). Secondary antibodies used were donkey anti-goat IgG AlexaFluor-594 (Invitrogen, A-11058, 1:250 dilution) and donkey anti-goat IgG AlexaFluor-488 (Invitrogen, A-11055, 1:250 dilution) for experiments involving co-staining for goat primary antibodies. Secondary antibodies were as follows: rabbit anti-goat-HRP (Jackson ImmunoResearch, 305-035-003, 1:10 000), goat anti-rabbit-HRP (Jackson Immuno120 Research, 111-035-144, 1:3000), donkey anti-sheep-HRP (Ab cam, ab6900, 1:5000), and sheep anti-mouse-HRP (Amersham GE, NA931, 1:2500).

### Histology

Hearts from 9-week-old control and mutant mice were carefully dissected, flushed with ice-cold PBS, and then fixed for 48 hours at 4 °C in a 4% paraformaldehyde solution (Sigma-Aldrich). Following fixation, the hearts were processed for paraffin-embedded sections and subjected to analysis through H&E staining, following the manufacturer’s protocol provided by Sigma-Aldrich.

### Immunofluorescence

Hearts from 9 weeks old control and mutant mice were dissected out and placed in ice-cold PBS, followed by freezing in OCT (Fisher) within the gas phase of liquid nitrogen. Cryosections of 7 μm thickness were mounted onto Superfrost Plus glass slides (Fisher Scientific), air-dried, and fixed for 10 minutes in ice-cold 4% paraformaldehyde (Sigma-Aldrich). Subsequently, sections were permeabilized in 1% Triton-X100 (Sigma- Aldrich) in PBS for 10 min, then blocked in 10% normal goat serum (Invitrogen) in PBS-T (PBS+0.1% Tween-20) for 30 min at room temperature (RT). Sections were then incubated overnight at 4°C in primary antibody diluted in blocking buffer, rinsed in PBS, then subsequently incubated for 60 min at RT in secondary antibody diluted in blocking buffer. Slides were mounted in VectaShield+DAPI (Vector Laboratories) or counterstained with DAPI and mounted in ProLog Gold (Invitrogen) priori to visualizing fluorescence. Primary antibodies were as follows: goat polyclonal anti-TBX3 (Santa Cruz, sc-31656, 1:250 dilution) and goat polyclonal anti-TBX5 (Santa Cruz, sc-17866, 1:250 dilution). Secondary antibodies used were donkey anti-goat IgG AlexaFluor-594 (Invitrogen, A-11058, 1:250 dilution) and donkey anti-goat IgG AlexaFluor-488 (Invitrogen, A-11055, 1:250 dilution) for experiments involving co-staining for goat primary antibodies. A secondary antibody-only control was employed in each case to ensure the specificity of immunostaining.

#### Statistics

The numbers of independent experiments are specified in the relevant figure legends. Quantitative data are presented as mean ± SD. Statistical analysis was performed using R version 3.6.2. or the GraphPad Prism statistical package version 10.2.1 (GraphPad Software, Boston, MA, USA). For all data sets, except qRT-PCR data, normality was assessed using Shapiro-Wilk normality test. If the data followed a normal distribution, the two-sample Welch t-test was performed for comparison. If normal distribution was not present, the non-parametric Mann-Whitney U test or Kruskal-Wallis H test were used, as indicated in the text. Statistical significance was assumed if P reached a value ≤ 0.05.

**For qRT-PCR data**, statistical analysis was performed using R version 4.2.0. Normality within each experimental group was assessed using the Shapiro-Wilk test. Between-group comparisons were conducted using Welch t-test, and multiple comparisons were corrected using the Benjamini & Hochberg method to control the false discovery rate (FDR) (71). If a significant difference was detected between two groups (t-test FDR < 0.05) but normality was rejected in any of the compared groups (Shapiro-Wilk P < 0.05), a non-parametric Wilcoxon rank-sum test was used for verification. A significant group-mean difference was confirmed at one-tailed Wilcoxon P≤0.05 (detailed in Supplementary Data Set I).

## Supplementary Materials

Supplementary Materials include Expended Method Section, Supplementary Figures 1 to 6 as well as Supplementary Data Set I are available online at eLife.

## Contributions

IPM and OBT conceived and designed the study; OBT, KAT, BL, MTB, ZK, ER, BL, ES, MG, KMS, and DEA performed wet lab experiments; OBT, KAT, BL, MTB, XHY, MG, KMS, IRE, and IPM participated in data analysis. IPM directed the project; OBT and IPM drafted the manuscript, with XHY providing revisions to the statistical analysis. OBT and IPM serve as co-corresponding authors. All authors contributed to and approved the final manuscript.

## Nonstandard Abbreviations and Acronyms

ACUP: animal care and use protocol
AER: atrial effective refractory period
A-H: atrio-hisian interval
APD50, 80: action potential duration at 50 and 80% of repolarization
AVB: atrioventricular bundle
AVERP: atrioventricular nodal effective refractory period
AVN: atrioventricular node
BBs: bundle branches
*Cacna1d/Cav1.3*: calcium channel, voltage-dependent, L type, alpha 1D subunit
*Cacna1g/Cav3.1d*: calcium channel, voltage-dependent, T type, alpha 1G subunit
*Cacna1h/Cav3.2*: calcium channel, voltage-dependent, T type, alpha 1H subunit
CCS: cardiac conduction system
Chr5: chromosome 5
*Col1a1*: collagen, type I, alpha 1
CREs: *cis*-regulatory elements
CRISPR: clustered regularly interspaced short palindromic repeats
EP studies: electrophysiology studies
ECG: electrocardiography
*Eyfp*: enhanced yellow fluorescent protein
FACS: fluorescent-activated cell sorting
FDR: False Discovery Rate
FS: fractional shortening
*Gja1/Cx43*: gap junction protein, alpha 1
*Gja5/Cx40*: gap junction protein, alpha 5
*Gjc1/Cx45*: gap junction protein, gamma 1
*Gjd3/Cx30.2*: gap junction protein, delta 3
GRN: gene regulatory network
GWAS: genome wide association studies
*Hcn1*: hyperpolarization activated cyclic nucleotide gated potassium channel 1
*Hcn4*: hyperpolarization-activated, cyclic nucleotide-gated K+ 4
Hd: his-duration
HOS: Holt-Oram syndrome
H-V: hisio-ventricular interval
IACUC: institutional animal care and use committee
Kcne1/MinK: potassium voltage-gated channel, Isk-related subfamily, member 1
*Kcnj2/Kir2.1*: potassium inwardly-rectifying channel, subfamily J, member 2
*Kcnj3/Kir3.1*: potassium inwardly-rectifying channel, subfamily J, member 3
*Kcnj4/*IRK3: potassium inwardly-rectifying channel, subfamily J, member 4
*Kcnj12/Kir2.2*: potassium inwardly-rectifying channel, subfamily J, member 12
*Kcnk3/Task-1*: potassium channel, subfamily K, member 3
lssDNA donor: long single-stranded DNA donor
LVEF: left ventricular ejection fraction
OAP: optical action potential
OE: overexpression
OMIM: online Mendelian inheritance in man
QRS: QRS complex
QT: QT-interval duration
PAM sequence: protospacer-adjacent motif sequence
*Postn*: periostin, osteoblast specific factor
PR: PR-interval duration
RFLP: Restriction fragment length polymorphism
*RR*: RR-interval duration
*Ryr2*: ryanodine receptor 2, cardiac
SAN: sinoatrial node
*Scn5a*/*Nav1.5*: sodium channel, voltage-gated, type V, alpha
sgRNA: single guide RNA
Smpx: small muscle protein, X-linked
*Tbx3*: T-box 3
*Tbx5*: T-box 5
TM: tamoxifen
VCS: ventricular conduction system
VERP: ventricular effective refractory period
VT: ventricular tachycardia

