## Supplementary material and figures for "Coordinated *Tbx3/Tbx5* transcriptional control of the adult ventricular conduction system"

Short title: ***Tbx3/Tbx5* patterns the cardiac conduction system**

Key words: *Tbx3, Tbx5,* T-box transcriptional factors, cardiac conduction system (CCS), ventricular conduction system (VCS), *Tbx3*:*Tbx5* double-conditional mouse line, *Tbx3*:*Tbx5*-deficient mice*,* reprogramming of VCS, heart rhythm, arrhythmia, heart patterning

*Please send correspondence to:
Ivan P. Moskowitz, M.D., Ph.D.
Departments of Pediatrics, Pathology, and Human Genetics
The University of Chicago
900 East 57th Street, KCBD Room 5102
Chicago, Illinois 60637
Phone: 773/834-0462


Ozanna Burnicka-Turek, Ph.D.
Departments of Pediatrics, Pathology, and Human Genetics
The University of Chicago
900 East 57th Street, KCBD Room 5240, LB1
Chicago, Illinois 60637
Phone: 773/702-2486


**This PDF file includes:**

Supplementary Data regarding validation of the newly generated *Tbx5* floxed allele

Supplementary Figures 1 to 6

**Supplementary Data**

*Validation of the newly generated Tbx5 floxed allele*

The *Tbx3:Tbx5* double-floxed mouse line (*Tbx3$^{fl/fl}$:Tbx5$^{fl/fl}$*) (see Methods section for details) was engineered by generating a novel *Tbx5* floxed allele in the background of a previously published *Tbx3* floxed allele (26). We utilized single *Tbx3* (26) and the novel *Tbx5* floxed mouse lines (*Tbx3$^{fl/fl}$* and *Tbx5$^{fl/fl}$*, respectively*)* to serve as controls in our studies (Figure 1A and Supplementary Figure 1 and 2). Although the newly engineered *Tbx5* floxed allele was designed to mirror a previously published allele (20), the location of the flox sites was altered slightly, requiring validation.

To verify the ability of the newly generated *Tbx5* floxed allele to recombine into the *Tbx5* KO (null) allele, we conducted three rounds of breeding and utilized a PCR assay to distinguish the recombined *Tbx5* KO (null) allele (*Tbx5$^{-}$*) from the unrecombined floxed allele (*Tbx5$^{fl}$*) (Supplementary Figure 2A and B). First, we bred newly generated homozygous *Tbx5* floxed males (*Tbx5$^{fl/fl}$*) with homozygous *Mef2C$^{Cre/Cre}$* females, which express *Cre* recombinase at the zygote stage (72-74), to produce germline *Tbx5$^{+/-}$:Mef2$^{Cre/+}$* double heterozygous mice. PCR analysis showed that 100% of the generated offspring were *Tbx5$^{+/-}$:Mef2C$^{Cre/+}$* double heterozygous mice (Supplementary Figure 2B). This result confirmed complete recombination of the new *Tbx5* floxed allele into the *Tbx5* null allele in the presence of the *Mef2C$^{Cre}$* allele. Next, we backcrossed *Tbx5$^{+/-}$:Mef2C$^{Cre/-}$* double heterozygote mice with wild-type CD1 IGS mice (Charles River Lab, USA) to obtain a germline *Tbx5$^{+/-}$* heterozygous mouse line (Supplementary Figure 2B). Subsequently, we bred *Tbx5$^{+/-}$* mice to generate germline *Tbx5* null mice (Tbx5$^{-/-}$). Among 230 offspring (generated in 50 litters), we observed 113 wild-type pups (*Tbx5$^{+/+}$*) and 116 *Tbx5* heterozygous pups (Tbx5$^{+/-}$), but no null newborn pups (*Tbx5$^{-/-}$*) were obtained (Supplementary Figure 2B). These results align with previous breeding outcomes indicating complete embryonic lethality of *Tbx5* null mice (20).

To further confirm the ability of the newly generated *Tbx5* floxed allele to produce germline *Tbx5* null embryos (*Tbx5$^{-/-}$*), we bred *Tbx5* heterozygous mice and collected embryos at *E9* to *E10*. Genotyping analysis of 89 collected embryos from 10 timed pregnancies, revealed the presence of 24 wild-type (*Tbx5$^{+/+}$*), 42 heterozygous (*Tbx5$^{+/-}$*), and 22 null (*Tbx5*$^{-/-}$) embryos (Supplementary Figure 2C). Supported by Chi-square analysis, these results collectively verified that the newly generated *Tbx5* floxed allele could recombine to form the null allele, mimicking the embryonic lethality observed with the previously generated allele (20).

To confirm that the newly engineered *Tbx5* conditional allele can generate VCS-specific *Tbx5*-deficient mice with the same phenotype as VCS-specific *Tbx5* knockout mice obtained from the previously generated allele (1, 11, 20), we crossed the newly generated *Tbx5* floxed mouse line (*Tbx5*$^{fl/fl}$) with a VCS-specific tamoxifen-inducible Cre transgenic mouse line (*MinK*$^{CreERT2}$ *[Tg(RP23-276I20-MinKCreERT2)]* (8)) (Figure 1A). The same tamoxifen regimen was administered as in the previous study (1, 11). We confirmed VCS-specific genetic deletion of *Tbx5* via immunofluorescence (IF) (Figure 1B) and qPCR (Figure 1C). The resulting VCS-specific *Tbx5*-deficient mice exhibited slowed conduction, indicated by increased PR and QRS intervals (Figure 2A, and 2E versus 2B), as well as episodes of ventricular tachycardia (VT) resulting in sudden death (Figure 2G and H), consistent with the previously reported phenotype (1, 11). Additionally, we confirmed the absence of histological and structural abnormalities in these mice, aligning with previous findings (Figures 3A, 3F versus 3B, and 3K versus 3G, respectively)(1, 11).

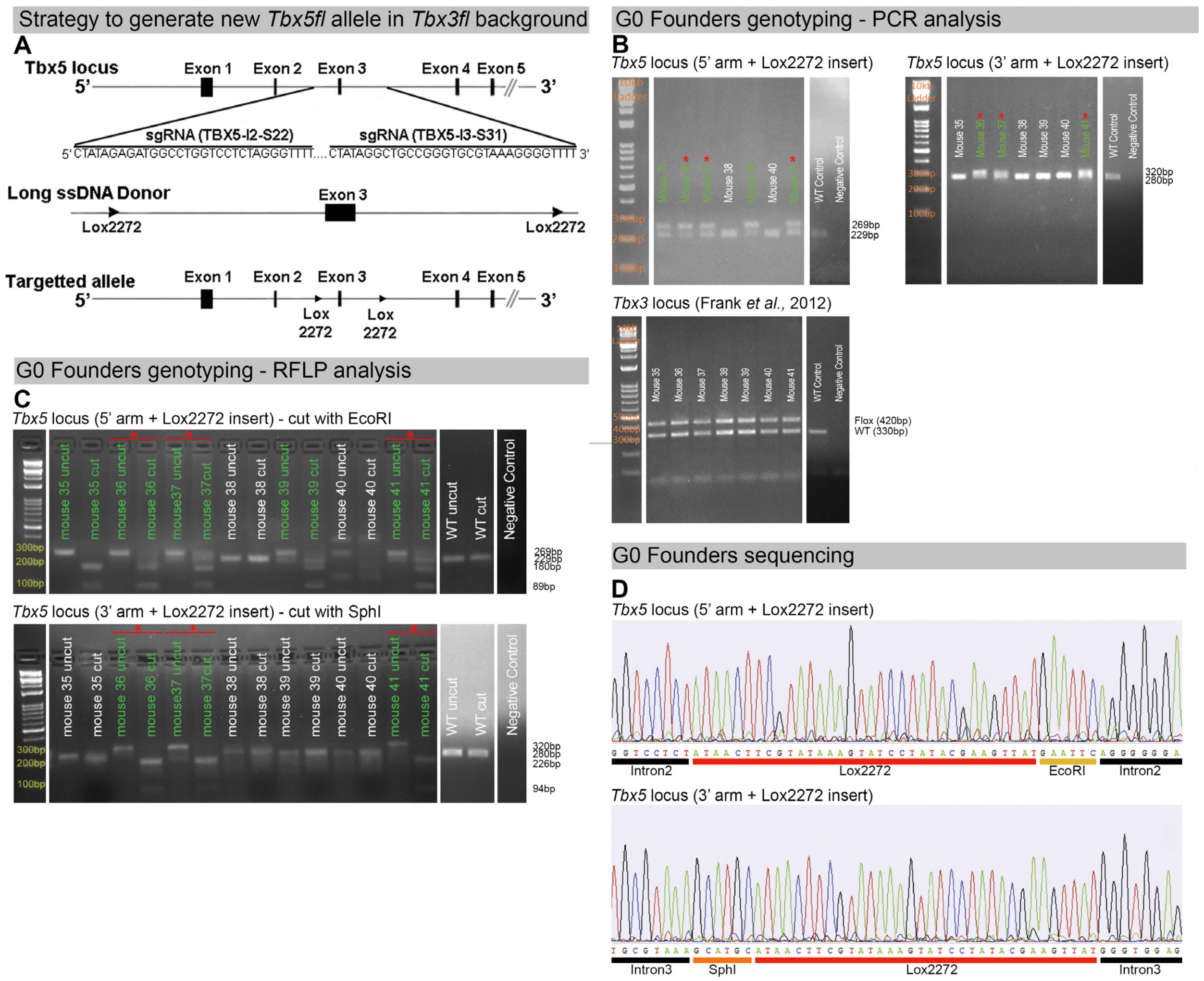


**Supplementary Figure 1. Generation of a novel *Tbx5* floxed allele in a *Tbx3* floxed background. (A)** Strategy to generate a new *Tbx5* floxed allele using the CRISPR/Cas9 system with long single-strand DNA donor (lssDNA donor) composed of the targeted exon 3 flanked by two lox2272 sites. The schematic shows the Cas9/sgRNA-targeting sites in the 2$^{nd}$ and 3$^{rd}$ introns of murine *Tbx5*, the lssDNA donor, and the targeted allele.**(B)** PCR analysis of *Tbx5* and *Tbx3* loci in G0 founders. Mice #35 to #41 are shown. **Upper left panel:** PCR genotyping of the *Tbx5* locus using primers specific for the 5' arm of the lssDNA donor with the lox2272 insertion. **Upper right panel:** PCR genotyping of the *Tbx5* locus using primers specific for the 3' arm of the lssDNA donor with the lox2272 insertion.

Green labels indicate founders positive for lox2272 insertion in one arm. Red asterisks indicate founders positive for lox2272 insertion in both arms. Expected fragment sizes: for the 5' arm with lox2272 insertion, positive band = 269 bp, WT = 229 bp; for the 3' arm with insertion, positive band = 320 bp, WT = 280 bp. **Lower panel:** PCR genotyping of the *Tbx3* locus using primers indicating *Tbx3* floxed allele. Expected fragment sizes: *Tbx3* floxed allele = 420 bp, Tbx3 WT allele = 330 bp. **(C)** RFLP analysis of the *Tbx5* locus in G0 founders. Mice #35 to #41 are shown. **Upper panel:** PCR products amplified using primers specific for the 5' arm of the lssDNA donor were digested with *EcoRI* to detect the insertion of the 5' lox2272. WT amplicon = 229 bp; lox2272 positive amplicon = 269 bp. The WT amplicon lacks the *EcoRI* site. With the 5’ lox2272 insertion, the digestion products are 180 bp and 89 bp. **Lower panel:** PCR products amplified using primers specific for the 3' arm of the lssDNA donor were digested with *SphI* to detect the insertion of the 3' lox2272. WT amplicon = 280 bp; lox2272 positive amplicon = 320 bp. The WT amplicon lacks the *SphI* site. With the 3’ lox2272 insertion, the digestion products are 226 bp and 94 bp. WT controls are shown in the last lanes. Green labels indicate founders positive for lox2272 insertion in one arm. Red asterisks indicate founders positive for lox2272 insertion in both arms. **(D)** Example of Sanger sequencing results from a representative G0 founder previously identified as positive for 5' and 3' lox2272 sites insertion by PCR and RFLP analysis. The sequencing confirmed the precise integration of lox2272 sites in both arms, resulting in exon 3 being flanked by two lox2272 sites.

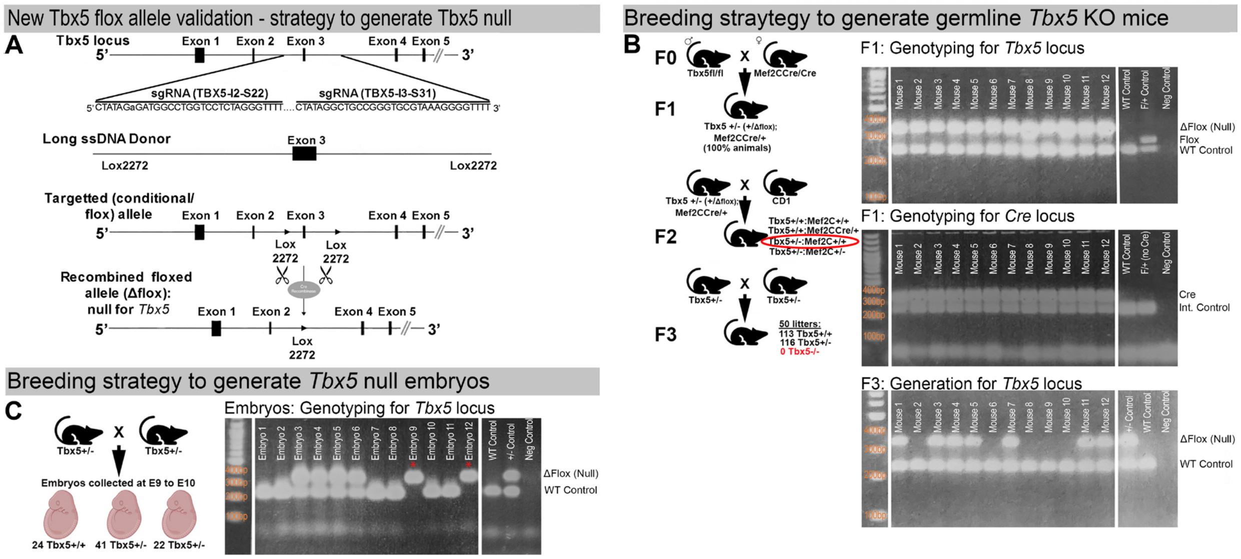


**Supplementary Figure 2. Validation of the newly generated *Tbx5* floxed allele. (A)** Strategy to generate a*Tbx5* null allele through recombination of the newly generated *Tbx5* floxed allele in the presence of the *Cre* recombinase. **(B)** Strategy to generate germline *Tbx5* null mice in presence of *Mef2C*$^{Cre}$ allele. To verify the ability of the newly generated *Tbx5* floxed allele to recombine into the *Tbx5* KO (null) allele, three rounds of breeding were conducted **(left panel)**, and a PCR assay **(right panels)** was utilized to distinguish the recombined *Tbx5* null allele (*Tbx5*$^{-}$) from the unrecombined floxed allele (*Tbx5*$^{fl}$). Initially, homozygous *Tbx5* floxed males (*Tbx5*$^{fl/fl}$) were bred with homozygous *Mef2C*$^{Cre/Cre}$ females, which express *Cre* recombinase at the zygote stage, to produce germline *Tbx5*$^{+/-}$*:Mef2*$^{Cre/+}$ double heterozygous mice. PCR analysis confirmed that 100% of the offspring were *Tbx5*$^{+/-}$*:Mef2C*$^{Cre/+}$ double heterozygous mice **(right upper and middle panels)**, demonstrating complete recombination of the *Tbx5* floxed allele into the *Tbx5* null allele in the presence of the *Mef2C*$^{Cre}$ allele. Subsequently, *Tbx5*$^{+/-}$*:Mef2C*$^{Cre/+}$ double heterozygous mice were backcrossed with wild-type CD1 IGS mice to obtain a germline *Tbx5*$^{+/-}$ heterozygous mouse line. *Tbx5*$^{+/-}$ mice were then bred to generate germline *Tbx5* null mice (T*bx5*$^{-/-}$). Among 230 offspring (from 50 litters), 113 wild-type pups (*Tbx5*$^{+/+}$) and 116 *Tbx5* heterozygous pups (*Tbx5*$^{+/-}$) were received, but no null pups (*Tbx5*$^{-/-}$) were obtained. **(Right lower panel)** Example of genotyping results showing absence of *Tbx5* null mice in newborn offspring generated by Tbx5$^{+/-}$ mice. Expected fragment sizes: *Tbx5 Δflox (null)* allele = 366 bp, *Tbx5 flox* allele = 269 bp, Tbx5 *WT* allele = 229 bp **(C)** Breeding strategy to generate *E9.0-E10.0* germline*Tbx5* null embryos. **(Left panel)** To generate germline *Tbx5* null embryos (*Tbx5*$^{-/-}$), *Tbx5* heterozygous mice (*Tbx5*$^{+/-}$) were bred, and embryos were collected at *E9* to *E10*. Genotyping analysis of 89 collected embryos from 10 timed pregnancies revealed the presence of 24 wild-type (*Tbx5*$^{+/+}$), 42 heterozygous (*Tbx5*$^{+/-}$), and 22 null (*Tbx5*$^{-/-}$) embryos. These results confirmed that the newly generated *Tbx5* floxed allele could recombine to form the null allele, mimicking the embryonic lethality observed with the previously generated allele (20). **(Right panel)** Example of genotyping results showing the presence of E9.0 *Tbx5* null embryos collected from *Tbx5+/-* timed breedings. Expected fragment sizes: *Tbx5 Δflox (null)* allele = 366 bp, *Tbx5 WT* allele = 229 bp.

Target Gene

gene name (abbr.) *Tbx5*

Ensembl Gene ID: ENSMUSG00000018263

Ensembl transcript ID: Tbx5-001 ENSMUST00000018407.9

sequence (5'->3', EXON 3: UPPERCASE, intron: lowercase)

200 bp buffer

[mark nucleotide sequence encoding critical residues in color]

ggtggttgataattgtttcacaatgaagagtagtgtctgactgttttggtattggtttt
tagaaataacttccagtggaaagagaaaatgttaaaaggcttaattggttgccccaagt
caaagaaaggctgggcggtggtgctagataaggcaagagtctggaaagattgttaatat
agacactgagggggaaaactactttttttggtccttgagaagaaaaacgattttaaaat
tgaaaatttctacatagtggctcacatgtgttggaatcttagcacttaggagggtgagg
caggagggttgccaagagtttgagaccacttttatgcgctgcatagtgaattccaggcc
tgtctgggctgcagagcaagactttgtctccaatattaataacaacaataatattgaaa
atttttgtatgttcatatttagacccagtgatgacggacagcgatgtatttgcatagct
gtgtgttttgcatctgggtgacaatacaaaaatatcctagtggggtttctagatggat
cccctcccaatcatgtttcagctaaaaatggatccaaatt**agatggcctggtcctctag**
**gggg**gagctctgtggaggggaataattcccaggcctcaatctggtggaagagacagcag
gtactaaggactgagatgctgctggacgcaagctccaactcacctagggagctgtaggc
agatgccatgggtcgccctagactttccatcactccccagaagtcagagcgctctttct
ttaaccgcacagactttggtaggtacttggcctctgcatatagggggagtttgagaaa
ggattccccgctaagacccatcactgaaaattttctctttgtctatcaag**GGCATGGAA**
**GGAATCAAGGTGTTTCTTCATGAACGTGAACTGTGGCTGAAGTTCCACGAAGTGGGCAC**
**AGAGATGATCATCACCAAGGCAGGGAG**gtgagccagctcctgccacagaaggaggacag
tcattggagggaatttaaaacaaaacaaaacaaaacaaaaaacacggaagaggcttggg
agaaaaaaattagaatagagccaatattcaatcagggttcattctggctaaagagtcct
ccgcaggatggagacctaacccccaatgcttacctccgggtaggatgttcgttcccga
ttttataggaggattaaagagaaaaaggtggaggcggggagaaaaggtgaagaacgggg
agactgtagaaggactttgagtccttagacttggacttgctccctggcctccacccta
ttcccaccttctctgaccttggaatagttccagggtccccacccgatgattgccaaacc
cgggattgttgaatgtattcgctacctcgttcattgtttatagagagccggtaaagtct
ttgaaagctagtggcgggggtggggtggggtagctcttcagcgctctgtacacagacac
gaataaacacagccgatcagctattgcgctgggcccaaactgctcttcaaggccggcag
gagtttcagctagagaggctaggctaggtctggggttggagggtgacagaagagttcaa
attctgtgtcttcggtgtgcatttgtgtgtgccggcctggggaacgacctcatttcccc
agcggcctgagctacatgtgtccccacaatataagaggccaccagtctgcccagtgcag
gggtgcggaagagacagaggccagggcagtaggcccaa**gctgccgggtgcgtaaagggt**
**gg**agttgattttccagattccgcgggttctccagcccaagccgccctcaataaactga
caacggtgacgtttattatttcttgaaattaaacaatgtcggctcccccgctcagcctg
aggcccttagtgaaccagaaggcggtgggtctgtttatatcagacatctgaggttccct
ttcccatcctgcccaggagggaaagtcccagccggctgccttgtcgtcctcgctggagc
cgagaggatttaactataaaggcctgcactgtctctggtgc

**Supplementary Figure 3. WT sequence of the targeted region at the mouse *Tbx5* locus.** The partial intron 2 (black, lowercase letters), entire exon 3 (black, bold, uppercase letters), and partial intron 3 (black, lowercase letters) are shown. The sgRNA- targeting sequence at the 5’ of exon 3 (TBX5-I2-S22) is labelled in bold, light green, while the sgRNA- targeting sequence at the 3’ of exon 3 (TBX5-I2-S31) is labelled in bold, dark green. The cleavage sites at the sgRNA-targeting sequences are underlined. The protospacer-adjacent motif (PAM) sequences are labelled in bold, light blue.

**Long ssDNA donor dsDNA template**

GGTACCatatcggatcccTAATACGACTCACTATAGgggtgacaatacaaaaatatcctagtgg
ggttttctagatggatcccctcccaatcatgtttcagctaaaaatggatccaaattagatggcc
tggtcctctATAACTTCGTATAAAGTATCCTATACGAAGTTATGAATTCagggggagctctgt
ggaggggaataattcccaggcctcaatctggtggaagagacagcaggtactaaggactgagatg
ctgctggacgcaagctccaactcacctagggagctgtaggcagatgccatgggtcgccctagac
tttccatcactccccagaagtcagagcgctctttctttaaccgcacagactttggtaggtactt
ggcctctgcatataggggggagtttgagaaaggattccccgctaagacccatcactgaaaattt
tctctttgtctatcaag**GGCATGGAAGGAATCAAGGTGTTTCTTCATGAACGTGAACTGTGGCT**
**GAAGTTCCACGAAGTGGGCACAGAGATGATCATCACCAAGGCAGGGAG**gtgagccagctcctgc
cacagaaggaggacagtcattggagggaatttaaaacaaaacaaaacaaaacaaaaaacacgga
agaggcttgggagaaaaaaattagaatagagccaatattcaatcagggttcattctggctaaag
agtcctccgcaggatggagacctaacccccaatgcttacctccgggtaggatgttcgttcccg
attttataggaggattaaagagaaaaaggtggaggcggggagaaaaggtgaagaacggggagac
tgtagaaggactttgagtccttagacttggacttgctccctggcctccacccctattcccacct
tctctgaccttggaatagttccagggtccccacccgatgattgccaaacccgggattgttgaat
gtattcgctacctcgttcattgtttatagagagccggtaaagtctttgaaagctagtggcgggg
gtggggtggggtagctcttcagcgctctgtacacagacacgaataaacacagccgatcagctat
tgcgctgggcccaaactgctcttcaaggccggcaggagtttcagctagagaggctaggctaggt
ctggggttggagggtgacagaagagttcaaattctgtgtcttcggtgtgcatttgtgtgtgccg
gcctggggaacgacctcatttccccagcggcctgagctacatgtgtccccacaatataagaggc
caccagtctgcccagtgcaggggtgcggaagagacagaggccagggcagtaggcccaagctgcc
gggtgcgtaaaGCATGCATAACTTCGTATAAAGTATCCTATACGAAGTTATgggtggagttgat
tttccagattccgcgggttctccagcccaagccgccctcaataaactgacaacggtgacgttt
attatttcttgaaattaaacaatgtcggctcccccgctcagcctgaggcccttagtgaaccaga
agGTCGAC

**KpnI SalI unique sites (**G GTAC C / G TCGA C)
**T7 promoter**
**Lox2272 sequence (ATAACTTCGTATAAAGTATCCTATACGAAGTTAT)**
***EcoRI* unique sites (G AATT C)**
***SphI* unique sites (**G CATG C)
**Left Homology Arm**
**Right Homology Arm**
**EXON 3**

**Supplementary Figure 4. Sequence of long ssDNA donor designed to target exon 3 of the mouse *Tbx5* locus.** It is composed of targeted exon 3 flanked by two lox2272 sites, along with 100/150 bp 5'/3' homology arms. The partial intron 2 (black, lowercase letters), entire exon 3 (bold, black, uppercase letters), partial intron 3 (black, lowercase letters), as well as 5’ (purple, lowercase letters) and 3’ (pink, lowercase letters) homology arms are displayed. The lox2272 sites, indicated in bold red, have been inserted at the cleavage sites of the sgRNA-binding sequences. Unique restriction enzymes sites used for RFLP analysis are shown as bold yellow for *EcoRI* and bold navy blue for *SphI*. T7 promoter is labeled in bold orange.

Ggtggttgataattgtttcacaatgaagagtagtgtctgactgttttggtattggttttaga
aataacttccagtggaaagagaaaatgttaaaaggcttaattggttgccccaagtcaaagaaa
ggctgggcggtggtgctagataaggcaagagtctggaaagattgttaatatagacactgaggg
ggaaaactactttttttggtccttgagaagaaaaacgattttaaattgaaaatttctacata
gtggctcacatgtgttggaatcttagcacttaggagggtgaggcaggagggttgccaagagtt
tgagaccacttttatgcgctgcatagtgaattccaggcctgtctgggctgcagagcaagactt
tgtctccaatattaataacaacaataatattgaaaatttttgtatgttcatatttagacccag
tgatgacggacagcgatgtatttgcatagctgtgtgttttgcatctgggtgacaatacaaaaa
tatcctagtggggtttctagatggatcccctcccaatcatgtttcagctaaaaatggatcca
aattagatggcctggtcctct**ATAACTTCGTATAAAGTATCCTATACGAAGTTAT**GAATTCag
gggggagctctgtggaggggaataattcccaggcctcaatctggtggaagagacagcaggtac
taaggactgagatgctgctggacgcaagctccaactcacctagggagctgtaggcagatgcca
tgggtcgccctagactttccatcactccccagaagtcagagcgctctttctttaaccgcacag
actttggtaggtacttggcctctgcatatagggggagtttgagaaaggattccccgctaaga
cccatcactgaaaatttctctttgtctatcaag**GGCATGGAAGGAATCAAGGTGTTTCTTCA**
**TGAACGTGAACTGTGGCTGAAGTTCCACGAAGTGGGCACAGAGATGATCATCACCAAGGCAGG**
**GAG**gtgagccagctcctgccacagaaggaggacagtcattggagggaatttaaaacaaaacaa
aacaaaacaaaaaacacggaagaggcttgggagaaaaaaattagaatagagccaatattcaat
cagggttcattctggctaaagagtcctccgcaggatggagacctaacccccaatgcttacct
ccgggtaggatgttcgttcccgattttataggaggattaaagagaaaaaggtggaggcgggga
gaaaaggtgaagaacggggagactgtagaaggactttgagtccttagacttggacttgctccc
tggcctccacccctattcccaccttctctgaccttggaatagttccagggtccccacccgatg
attgccaaacccgggattgttgaatgtattcgctacctcgttcattgtttatagagagccggt
aaagtctttgaaagctagtggcgggggtggggtggggtagctcttcagcgctctgtacacaga
cacgaataaacacagccgatcagctattgcgctgggcccaaactgctcttcaaggccggcagg
agtttcagctagagaggctaggctaggtctggggttggagggtgacagaagagttcaaattct
gtgtcttcggtgtgcatttgtgtgtgccggcctggggaacgacctcatttccccagcggcctg
agctacatgtgtccccacaatataagaggccaccagtctgcccagtgcaggggtgcggaagag
acagaggccagggcagtaggcccaagctgccgggtgcgtaaa**GCATGCATAACTTCGTATAAA**
**GTATCCTATACGAAGTTAT**gggtggagttgattttccagattccgcgggttctccagcccaa
gccgccctcaataaactgacaacggtgacgtttattatttcttgaaattaaacaatgtcggct
cccccgctcagcctgaggcccttagtgaaccagaaggcggtgggtctgtttatatcagacatc
tgaggttccctttcccatcctgcccaggagggaaagtcccagccggctgccttgtcgtcctcg
ctggagccgagaggatttaactataaaaggcctgcactgtctctggtgc

**Lox2272 sequence (ATAACTTCGTATAAAGTATCCTATACGAAGTTAT)**
***EcoRI* unique sites (G AATT C)**
***SphI* unique sites (**G CATG C)
**EXON 3**
introns

**Supplementary Figure 5. Sequence of mouse new *Tbx5* floxed allele generated using CRISPR/Cas9 system with lssDNA composed of targeted exon 3 flanked by two lox2272 sites.** The partial intron 2 (black, lowercase letters), entire exon 3 (bold, black, uppercase letters), and partial intron 3 (black, lowercase letters) are displayed. The lox2272 sites, indicated in bold red, have been inserted at the cleavage sites of the sgRNA-binding sequences. Unique restriction enzymes sites used for RFLP analysis are shown as bold yellow for *EcoRI* and bold navy blue for *SphI*. PCR primers specific for the 5' arm of the lssDNA donor with the lox2272 insertion (displayed as green box) as well as primers specific for the 3' arm of the lssDNA donor with the lox2272 insertion (displayed as pink box) are shown.

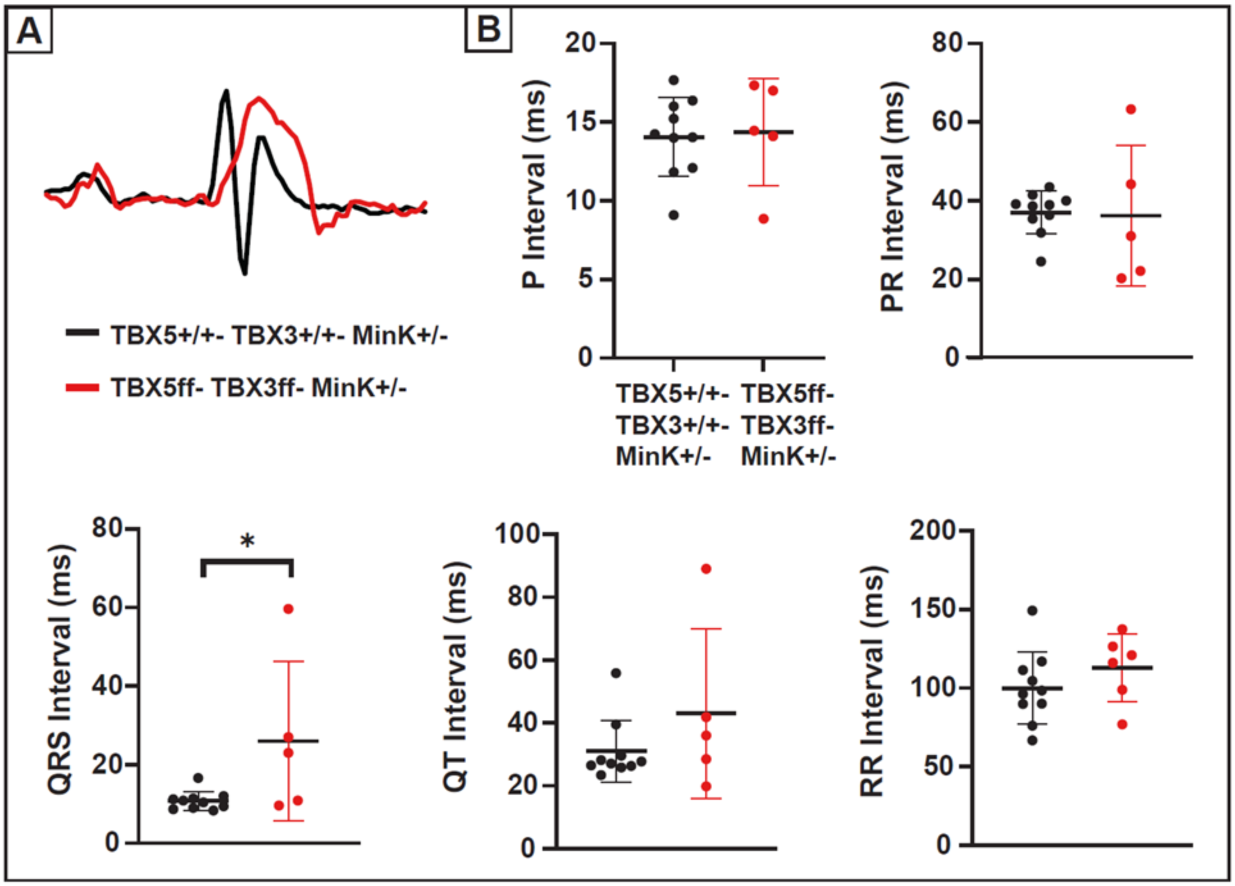


**Supplementary Figure 6. *Tbx3:Tbx5* double-conditional knockout mice exhibit QRS prolongation. (A)** Representative ECG traces recorded from *Tbx3:Tbx5* double-conditional knockout mice (red) and control littermates (black). (**B)** Quantification of P, PR, QRS, QT, and RR intervals. *$P$<0.05 denotes significance.
